# TASC: A transcriptome-driven machine learning classifier to explore molecular heterogeneity and relapse-associated programs in T-cell Acute Lymphoblastic Leukemia

**DOI:** 10.64898/2025.12.23.696007

**Authors:** Cinzia Benetti, Omar Almolla, Giovanni Gambi, Annamaria Massa, Iannis Aifantis, Aristotelis Tsirigos, Francesco Boccalatte

## Abstract

T-cell acute lymphoblastic leukemia is a biologically heterogeneous malignancy characterized by diverse transcriptional and genomic alterations. Recent studies have defined a set of recurrent molecular subtypes associated with distinct differentiation stages and clinical outcomes. However, no unified framework currently exists for assigning these subtypes in a standardized and accessible manner. Existing approaches often rely on mutation or fusion detection and may overlook broader transcriptional programs. The lack of a comprehensive, transcriptome-based classification tool has hampered the use of subtype-specific insights in both research and clinical settings. Here, we present a machine learning–based classifier trained on transcriptomic data to predict previously defined multi-omic subtypes of T-cell acute lymphoblastic leukemia. The model accurately assigns subtype identity across patient samples and cell lines, and provides a practical tool for standardized molecular stratification, supporting future integration into diagnostic and translational workflows, as demonstrated by its ability to reveal subtype-specific patterns of relapse.

## Introduction

T cell acute lymphoblastic leukemia (T-ALL) is an aggressive hematological malignancy affecting the T cell compartment, particularly frequent in the pediatric age(*1*).

According to the most recent World Health Organization classification of hematological tumors (WHO-HEM5) in 2022(*2*, *3*), T-ALL is recognized as a single diagnostic entity, with the only exception of the early T-cell precursor acute lymphoblastic leukemia (ETP-ALL), which gained formal recognition based on its peculiar immunophenotypic profile(*4*). Nonetheless, T-ALL is a molecularly heterogeneous malignancy characterized by a wide spectrum of genomic alterations, leading to the aberrant expression of transcription factors and oncogenes(*5*). These events drive the activation of distinct downstream pathways, resulting in subtype-specific transcriptional programs often associated with defined stages of normal T-cell development(*5–10*).

To overcome the coarseness of the current stratification, the latest revision of the International Consensus Classification(*11*) added eight provisional entities for T-ALL based on the aberrant activation of specific transcription factor families (TAL1, TLX3, LMO2, BCL11B, SPI1 and HOXA being the most studied)(*12–19*). Despite substantial progress in uncovering the molecular heterogeneity of T-ALL, there remains no unified framework for its subtype classification. This lack of standardization continues to hinder systematic subtype representation, limiting both translational research efforts and the implementation of precision medicine strategies in clinical settings(*20*, *21*). Notably, T-ALL remains one of the least molecularly stratified hematologic malignancies, especially when compared to B-cell(*22*, *23*) or myeloid malignancies, where robust and widely adopted classification frameworks have already been established(*23*). Existing approaches have primarily relied on the detection of recurrent coding mutations through whole-exome sequencing (WES) and whole-genome sequencing (WGS)(*24*), the identification of gene fusions from bulk RNA sequencing (RNA-seq) data(*24*, *25*) or oncogene-driven gene expression approaches(*26*). Given the complex interplay of genetic and epigenetic factors influencing T-ALL heterogeneity, recent large-scale integrative studies have employed genomic, epigenomic, and transcriptomic profiling to achieve more refined disease stratification(*25*, *27*, *28*). These efforts culminated in a recent publication by Pölönen and colleagues, who identified 15 biologically and clinically distinct subtypes, each defined by unique genomic alterations, transcriptional programs, and differentiation states(*28*). These provisional subtypes differ from the diagnostic and prognostic standpoint, which underlines the need for a comprehensive characterization to recognize subtypes(*20*).

In this study, we introduce our T-cell Acute Lymphoblastic Leukemia Subtype Classifier (TASC), a Random Forest–based transcriptional classifier implemented as an accessible web application (Shiny App). Rather than serving solely as a subtype assignment tool, TASC was intentionally trained on well-defined subtypes and biologically relevant features, to enhance interpretability and downstream analytical utility, then tested and validated on a large cohort of public datasets (Supplementary Data 1). TASC provides a unified and user-friendly classification framework reliant on readily obtainable transcriptomic data, which proves to be useful in assessing other aspects beyond patient stratification, such as identifying intra-patient heterogeneity and relapse-driven pathways in T-ALL.

## Results

### Defining model features

To build a transcriptional classifier for T-ALL, we first aimed to identify predictive features capable of distinguishing each class. To this end, we analyzed transcriptomic data from the largest dataset available so far(*28*), containing 1309 samples annotated according to the 15 most recent and comprehensive subtypes(*28*). We selected features in a biology driven manner, prioritizing explainability, in light of the highly heterogenous nature of T-ALL. We therefore selected the 300 most variable genes (MVGs) from the original dataset(*28*), which were also used for subtype clustering(*28*), as effective descriptors of variance among individual groups (**Figure 1a**). While overexpression of only a few of MVGs could be attributed to individual classes, more complex subtype-defining expression patterns emerged, supporting MVGs’ utility in subtype discrimination (**Figure 1b**). To complement this, we calculated the differentially expressed genes (DEGs) in each subtype (**Supplementary Data 2**). Although MVGs and DEGs showed limited overlap (**Supplementary Figure 1a**), both sets provided clear transcriptomic-based discrimination of subtypes (**Figure 1b-c**).

**Figure 1:**
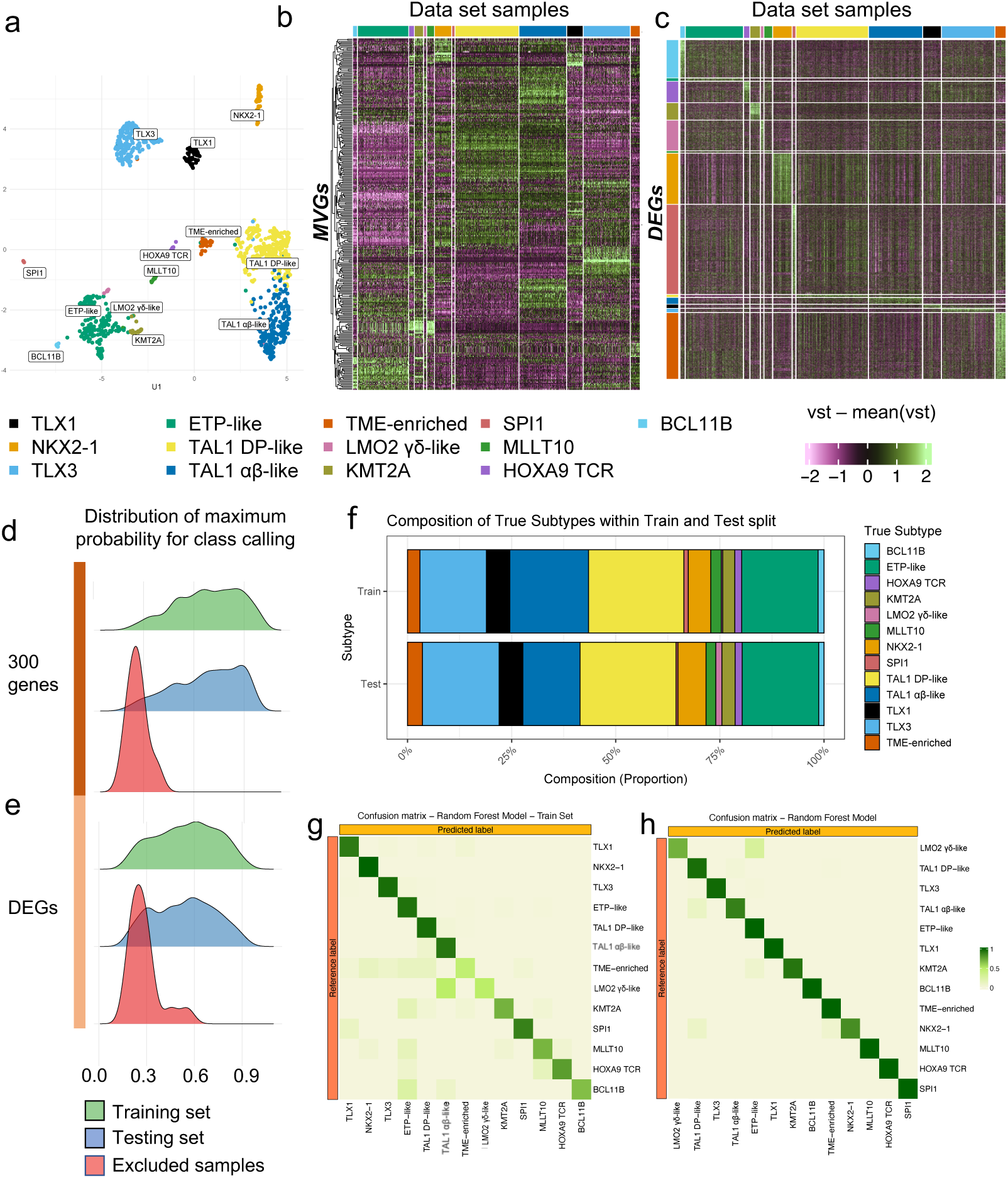
The 300 most variably expressed genes (MVGs) outperform class-specific DEGs in discriminating selected multi-omic subtypes. **a,** UMAP projection of ALL patient samples from 13 subtype groups (each >1% prevalence), generated using the top 300 most variable genes(*28*). Sample points are colored by subtype. **b,** Heatmap of variance-stabilized, mean-scaled expression of MVGs, organized by subtype (columns), with gene modules ranked by hierarchical clustering. **c**, Heatmap of variance-stabilized, mean-scaled expression of DEGs, where columns are split and colored by ground truth labels and rows are split and colored by DEGs specific to each subtype. **d**, Density distributions of maximum posterior probabilities RF classifier for each subtype, shown for MVG RF model, split into training data (green), held-out testing samples (blue), and samples from excluded subtypes (STAG/LMO2 - NKX2-5, red). **e**, same as **d** but referring to the DEGs RF model. **f**, Bar plot of dataset composition, comparing ground truth subtype proportions in the train and test set. **g,** Confusion matrix summarizing RF performance on the training set using the MVG-based model. Rows denote true subtype labels; columns are model predictions. Color intensity reflects per-cell classification accuracy. **h**, same as **g** but on the testing set

To evaluate the class-predictive potential of these gene sets, we trained two separate Random Forest (RF) models(*29*) using variance stabilizing transformation (vst) normalized expression values(*30*) as input features. The models were trained on a random subset comprising 63% of the full dataset, leveraging existing subtype annotations(*28*), with the remaining samples used for testing.

Subtypes represented by fewer than ten samples lacked sufficient representation to support reliable bootstrap sampling and model training, and their inclusion resulted in unstable predictions and reduced overall model performance (**Supplementary Table 1**). Consistent with recommendations from the literature(*31*, *32*), we re-analyzed the data after excluding these low-incidence subtypes, which led to improved performance metrics across both models (**Supplementary Data 3**). To rule out the possibility of data leakage stemming from the biologically informed feature selection, we performed a label permutation control: for each model, the subtype labels were randomly shuffled, and the models were re-trained on these randomized labels. Performance was then evaluated on the held-out testing set using p-ROC AUC, and compared to the AUC obtained with the true labels (**Supplementary Figure 1b**). The substantial drop in predictive performance upon label randomization demonstrates that the models learn genuine subtype-associated transcriptomic signals rather than artefactual correlations, providing a rigorous proof of the predictive value of the selected features (**Supplementary Data 4**). Between the two predictor sets, MVGs yielded superior model performance, with balanced accuracy increasing from 0.76 (range: 0.500 for LMO2 γδ-like to 0.971 for TLX3) to 0.94 (range: 0.750 for SPI1 to 0.998 for HOXA TCR) in the MVG RF model, and F1-scores rising from 0.577 (0 for LMO2 γδ-like to 0.918 for TLX3) to 0.915 (0.667 for SPI1 to 0.986 for NXK2-1) (**Supplementary Data 3**). Moreover, the distribution of maximum posterior probabilities per class of the MVG-based model displayed a higher maximum peak and a smoother, more gradually increasing slope, indicative of greater confidence and calibration in class assignment(*33*) (**Figure 1 d-e**).

We also assessed model calibration by calculating the Expected Calibration Error (ECE) and Brier score for each model. Both the MVG- and DEG-based models exhibited a tendency towards under-confidence, i.e., they predicted correct subtypes with lower-than-expected probabilities (**Supplementary Figure 1c**). Notably, the DEG-based model displayed higher ECE and Brier scores than the MVG-based model (ECE: DEG = 0.33, MVG = 0.28; Brier score: DEG train = 0.32, test = 0.33; MVG train = 0.21, test = 0.19), indicating that, while both models achieve good classification accuracy, the MVG model provides slightly better-calibrated probability estimates. Overall, the subtype composition of the train and test set is similar, with minimal difference in LMO2 and SPI1 composition (**Figure 1f**), which could explain the Brier score differences between train and test sets in the MVG model. This is also supported by confusion matrices for the training set (**Figure 1g**, **Supplementary Data 5**) and the testing set (**Figure 1h**) further illustrated the performance of the MVG-based model. While the LMO2 γδ-like class showed the lowest sensitivity, often misclassified as TAL1 αβ-like and ETP-like, it exhibited a high Precision Recall Area Under the Curve (PR-AUC) (**Supplementary Data 6**). This is consistent with the distinct probability distribution of this class in true LMO2 γδ-like compared to others, suggesting the potential for inclusion of additional cases based on prediction probabilities (**Supplementary Figure 1d**). In the two RF models (built using either MVGs or DEGs), samples belonging to excluded subtypes displayed lower maximum probabilities (median <0.3), confirming suboptimal classification for these samples compared to those from included subtypes (**Figure 1d-e**). Most of the excluded samples from NKX2-5 and STAG2/LMO2 subtypes were classified as ETP-like and TAL1 (**Supplementary Figure 1e**), potentially due to the higher relative abundance of the latter (**Supplementary Table 1**). However, the MVG-based model demonstrated a greater separation between recognized and unrecognized samples, supporting its utility for robust subtype classification. Therefore, we adopted the MVG-based RF model, hereafter referred to as TASC (T-ALL Subtype Classifier), as the primary method for subtype prediction.

### Performance of TASC in subtype classification

To further evaluate the performance of TASC relative to other machine learning approaches, we compared the two RF models with an XGBoost(*34*) classifier trained on MVGs. Among the three, TASC consistently achieved the highest overall performance, emerging as the only model to obtain admissible F1 scores and balanced accuracy in the LMO2 γδ-like subtype (**Figure 2** and **Supplementary Data 5**). While both MVG-based models (RF and XGBoost) displayed broadly comparable performance across the other subtypes, XGBoost achieved slightly higher accuracy. In contrast, TASC consistently delivered superior F1 scores, reflecting a more balanced trade-off between precision and recall. These metrics indicate that TASC offers competitive performance relative to other machine learning models, combining strong accuracy with improved balance between precision and recall.

**Figure 2:**
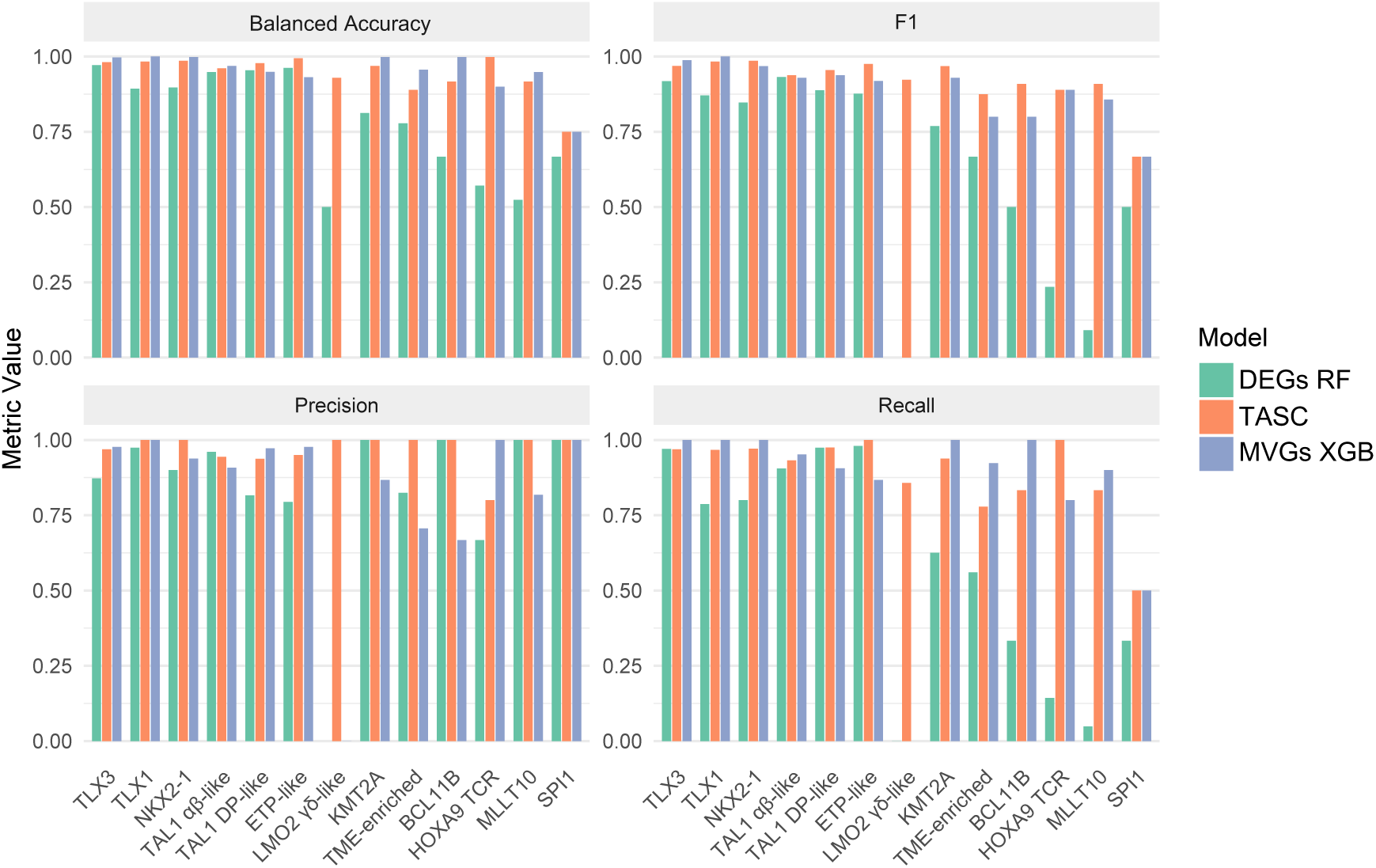
Evaluating TASC quantitative metrics. Bar plots of classification metrics (Balanced Accuracy, F1, Precision, and Recall) for each subtype in the test set. Each plot compares three models, each represented by a different color: TASC (orange), RF using DEGs (green), and XGBoost using MVGs (blue).

### Model explainability reveals novel subtype features

Global feature importance, assessed using cross-entropy loss, identified key class-defining oncogenes (such as *TLX3*, *TLX1*, and *HOXA9)* among the top 20 predictive features (**Figure 3a**), in line with established associations between subtype identity and oncogene expression(*5*). Notably, the top features also included genes implicated in T-cell development and T-ALL pathogenesis, such as *CD34* and *PROM1*(*35–37*) (**Figure 3a**), the latter encoding a component of the PI3K signaling pathway, which regulates tumor growth, cell survival, and drug resistance(*38*). To gain subtype-level interpretability, we applied SHapley Additive exPlanations (SHAP), a game-theoretic framework for explaining individual predictions of machine learning models(*39*). We computed local SHAP values for each class to evaluate the contribution of specific genes to subtype assignment. Clustering of SHAP values across subtypes reflected biologically meaningful class proximities, such as between *MLLT10* and *HOXA9 TCR*, or *ETP-like* and *BCL11B* (**Figure 3b-c**)(*28*). Interestingly, comparing clustering based on the expression of the RF model predictors with clustering derived from their SHAP importance revealed that feature importance captured aspects beyond expression dynamics across subtypes, as reflected by the differences in subtype clustering (**Figure 3b–c**). SHAP importance metrics more accurately capture the similarity between the LMO2 γδ-like and TAL1 subtypes, reflecting their known biological resemblance and non-mutually exclusive nature (*40*). SHAP rankings were also concordant with Gini-based importance measures (Pearson = 0.88), supporting their interpretability (**Figure 3d**). Overall, SHAP values partially reflected what was observed in the cross-entropy loss calculations, as *TLX1* and *TLX3* were important features for several classes. In the respective subtypes (e.g., TLX3), high feature values favored the model decision toward that class, while the opposite could be observed in other contexts (**Figure 3e-f**). Interestingly, in the ETP-like subtype, the most influential predictive features identified by SHAP were primarily negative contributors (**Figure 3e**), potentially accounting for the reduced specificity observed in this class (**Figure 1g**). Nevertheless, the small subset of genes acting as positive predictors revealed biologically relevant associations. Among them, *HOXA9* is a gene recurrently deregulated by enhancer hijacking in ETP-like cases(*28*), while *HPGDS*, a metabolic regulator(*41*), and *MN1*, a transcription factor implicated in stemness, adverse prognosis across malignancies, and regulation of *CD34* expression, emerge as putative markers of this subtype(*42–44*). To assess whether the relative scarcity of positive predictors among highly ranked features of ETP-like subtype increased susceptibility to false-positive classification in non–T-ALL samples, we applied the classifier to an independent cohort of 499 B-ALL cases (**Supplementary Data 7**). In the absence of probability thresholds, the majority of samples were assigned highest probability to the ETP-like subtype (**Supplementary Figure 2a**), indicating reduced subtype specificity. Therefore, we decided to calculate thresholds on the testing set to allow both the recovery of LMO2 γδ-like samples and the exclusion of the ETP-like false positive cases, introducing the “NA” fallout class.

**Figure 3:**
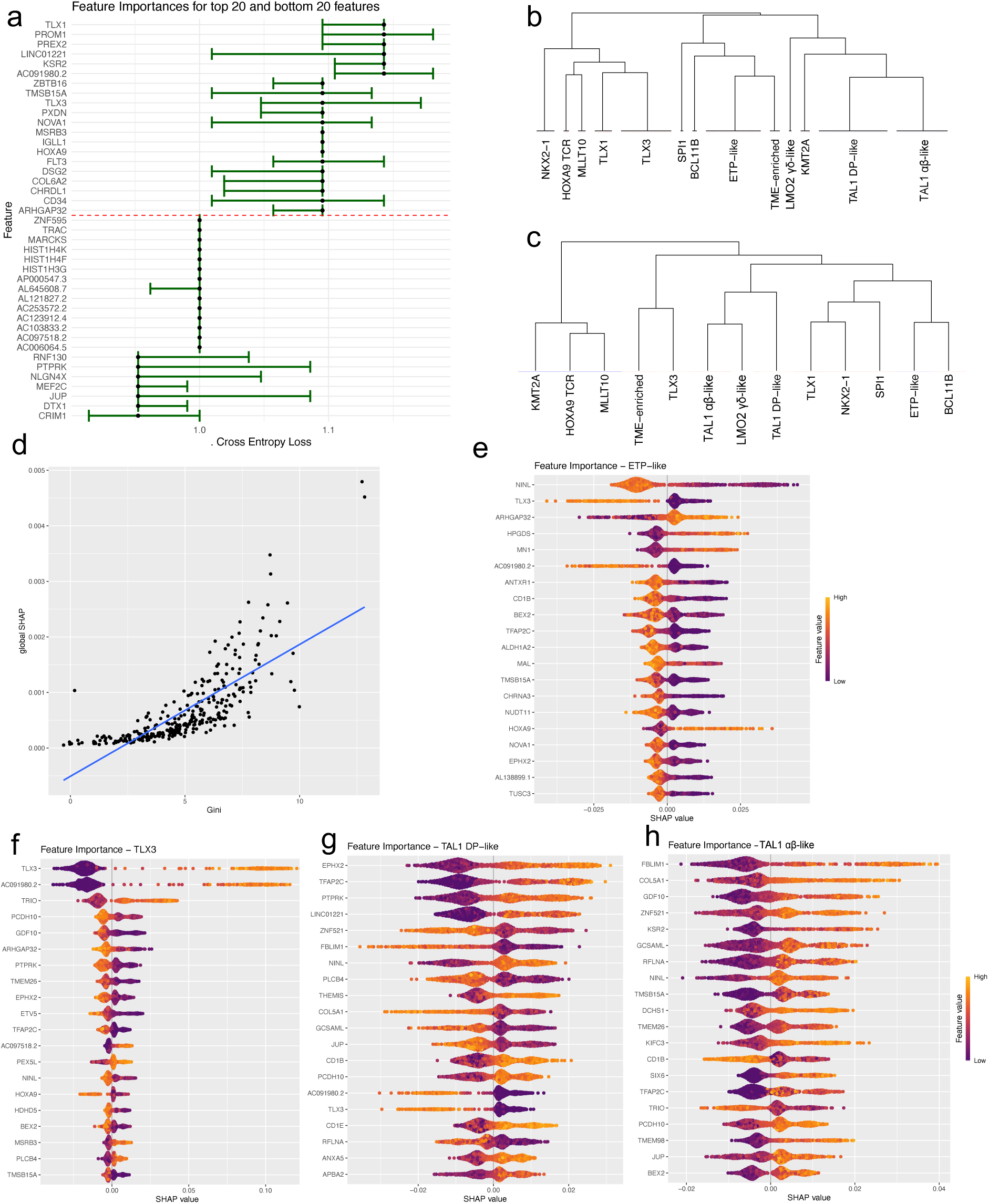
Model interpretability analysis. **a,** Forest plot displaying feature importance ranked by cross-entropy loss. The top 20 and bottom 20 ranked features (y-axis) are shown with their estimated effect on the model’s loss (x-axis) and 95% confidence intervals. **b,** Subtype-grouped dendrograms generated from variance-stabilized transformed expression profiles of MVGs, using Euclidean distance and complete linkage method for clustering. **c,** Dendrograms derived from median-scaled SHAP values using Pearson correlation distance and Ward D2 clustering, highlighting model-driven subtype similarity. **d**, Pearson correlation between mean SHAP values across classes and mean decrease in gini coefficient, highlighting **e-h,** Feature importance plots for ETP-like (**e**) TLX3 (**f**) TAL1 DP-like (**g**) and TAL1 αβ-like (**h**) subtypes. Each dot represents a sample SHAP value (x-axis) for each of the top 20 features (y-axis) ranked according to absolute mean SHAP value. Color represents vst gene expression for the corresponding feature, which determines if the feature is a positive (higher expression linked to higher SHAP values) or negative predictor (higher expression linked to lower SHAP values) for the model in the given class.

Thresholds were defined using ROC AUC analysis, including Youden’s index and a more stringent, clinically oriented threshold prioritizing high specificity (specificity and sensitivity >90%, with maximal specificity, **Supplementary Table 2**).

Application of these thresholds substantially improved classifier performance in the B-ALL cohort (**Supplementary Figure 2b–c**). Using Youden-maxima thresholds, over 75% of samples were reassigned to the NA class, with remaining predictions primarily corresponding to ETP-like, KMT2A, or dual ETP-like/KMT2A assignments (**Supplementary Figure 2b**). Notably, non-NA KMT2A and ETP-like/KMT2A cases were enriched for B-ALL samples harboring KMT2A alterations (**Supplementary Figure 2d**).

Under the more stringent clinical threshold, only two B-ALL samples remained unassigned, one classified as ETP-like and one as KMT2A, both carrying documented KMT2A alterations. Together, these findings indicate that, despite reduced specificity in the absence of thresholding, the classifier captures biologically meaningful transcriptional features consistent with known leukemia subtypes.

Local SHAP value analysis further highlights the model’s capacity to discriminate between transcriptionally similar subtypes. This is particularly evident in the distinction between the two TAL1-altered subtypes, TAL1 DP-like and TAL1 αβ-like, which, despite sharing a common oncogenic driver, differ markedly in differentiation state and transcriptional profiles. Notably, the most influential features vary substantially between these subtypes, with genes such as *ZNF521*, *FBLIM1*, and *CD1B* exhibiting opposite SHAP contributions (**Figure 3g-h**). These genes shift from being strong positive predictors in one subtype to negative predictors in the other, underscoring the model’s sensitivity to subtype-specific gene regulation and its ability to resolve fine-grained biological differences.

Overall, feature importance in TASC reveals subtype-specific biological patterns aligned with T-ALL biology.

### Robustness of subtype predictions under feature perturbation and batch effects

To evaluate the robustness of subtype predictions and to exclude potential technical confounding, we systematically perturbed model inputs at multiple levels, including feature identity, feature values, and batch structure.

First, we assessed sensitivity to feature identity by permuting genes in ranked sliding windows of 30 genes according to training-set importance. Disruption of top-ranked genes led to a non-linear reduction in test-set performance, with a marked drop in accuracy in the first 15 windows, whereas permutation of lower-ranked genes had minimal impact on mean accuracy (**Supplementary Figure 3a**). Probability-AUCs displayed overall logarithmic growth, with a smaller decrease in mean AUC (**Supplementary Figure 3b**). This behavior is inconsistent with diffuse feature contribution and instead indicates reliance on a constrained subset of subtype-informative transcriptional programs.

Next, we evaluated the impact of progressive gene imputation, excluding the top 30 SHAP-ranked features that drive the largest drop in accuracy when perturbed. Increasing levels of imputation resulted in a systematic reduction of predicted class probabilities (**Supplementary Figure 3c**), demonstrating that imputation affects the model’s confidence in a non-trivial manner. Importantly, in an independent external cohort of 37 bulk RNA-seq samples from the Single Cell Pediatric Cancer Atlas (**Supplementary Data 1**), sequential imputation up to approximately 40 genes did not substantially alter class assignments or predictive accuracy, with only a few first positives appearing at the highest imputation levels (**Supplementary Figure 3 d-f**). Together, these results suggest that while imputation modulates prediction probabilities, it does not compromise the model’s overall classification performance, which allowed us to extend our validation cohort and more easily adapt the model for general use, by allowing imputation on a maximum set of 30 genes on the whole train-test cohort, excluding top ranked features.

Finally, we examined the combined effects of batch and imputation on prediction probabilities by comparing maximum class call probabilities across batches, including on top of the 37 samples from the imputation validation cohort, 85 samples from two independent cohorts, annotated according to the most recent transcriptional subtype classifier, TALLSorts(*45*), which required imputation of 25 features. This revealed how despite strong batch effect among cohorts (**Supplementary Figure 3g**), no significant difference in linear regression was observed relative to the test set when batch effect is present alone (**Supplementary Table 3**), despite strong differences in subtype composition (**Supplementary Figure 3h**). In contrast, when both batch and imputation were present, the effect was strongly significant (**Supplementary Table 3**). These findings suggest that probability thresholds should be applied selectively upon imputation, as it affects probability distributions, particularly focusing on ETP-like and LMO2 driven subtypes, to reduce false positives and false negatives, while retaining overall predictive performance.

### Validation of TASC on external cohorts

To validate TASC robustness, we applied the model on the external dataset comprising 85 samples with oncogene level annotations, used to train the TALLSorts classifier(*45*). TASC predictions demonstrated high concordance with these ground truth labels, particularly for canonical subtypes such as TLX3, KMT2A, MLLT10, and TAL1. TLX1 showed near-perfect alignment, except for three TLX1 samples that were misclassified as either TLX3 (2) or TAL1 DP-like (1) (**Figure 4a**). Samples previously annotated as “diverse”, a category known to be enriched for ETP-ALL cases(*45*), were consistently reassigned to the ETP-like and BCL11B subtypes by TASC, further reflecting their shared molecular features and ETP-related biology.

**Figure 4:**
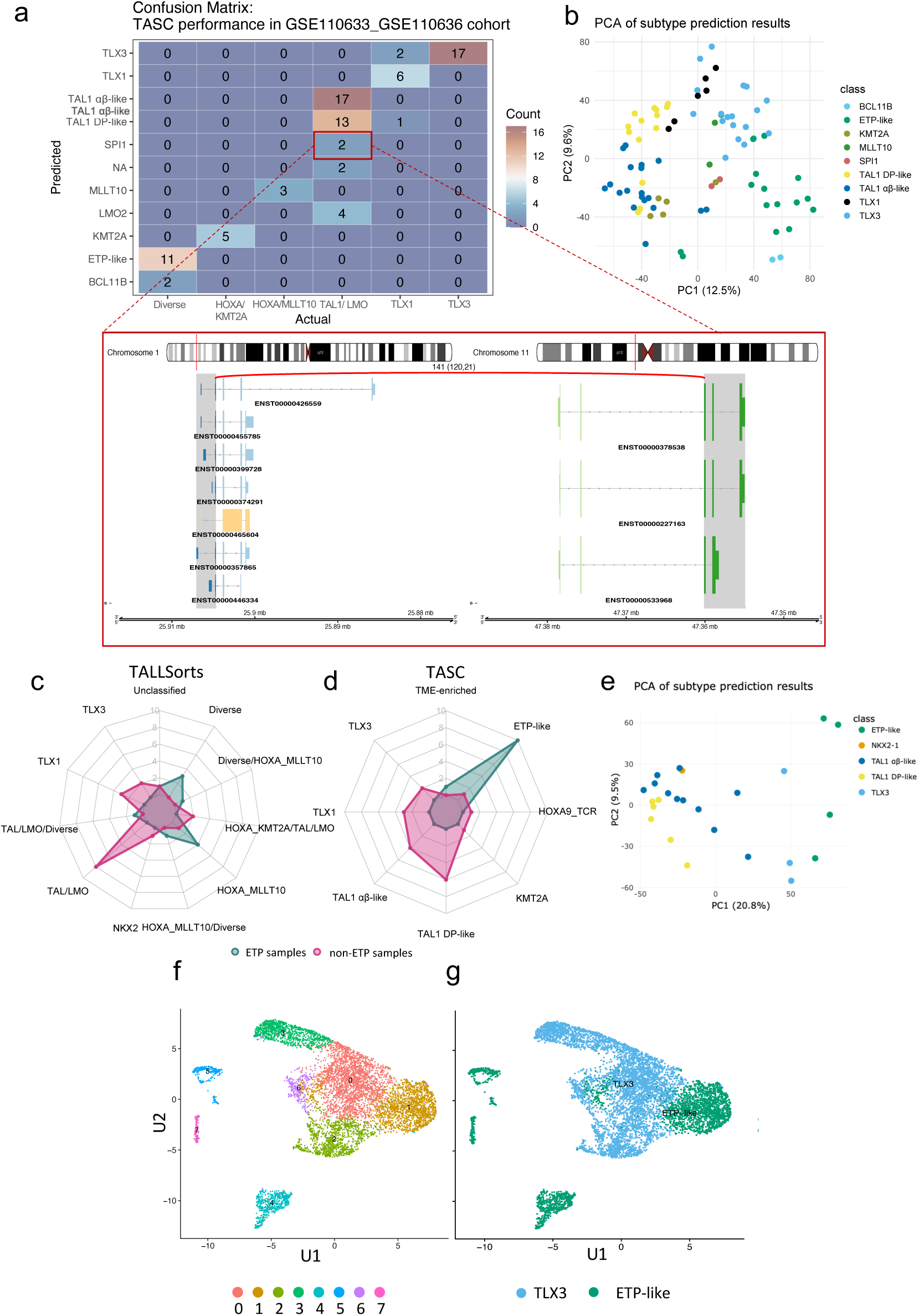
Robust validation and benchmarking across a diverse sample panel. **a,** Validation on 85 samples from two independent cohorts (GSE110633 and GSE110636)(*45*), using ground-truth labels from the TALLSorts training set as reference. Model predictions are shown alongside true labels. For two apparently misclassified SPI1 samples, a gene-fusion plot showcasing the *SPI1* fusion event detected by the STAR-fusion pipeline on raw RNA-seq data is included. **b,** Principal component analysis (PCA) on combined GSE110633 and GSE110636(*45*) cohorts, showing PC1 vs. PC2 with percentage variance explained. Each point represents a sample colored by subtype. **c,** Radar plot comparing the predictions for two groups of samples (ETP and non-ETP T-ALL) from the GSE243914(*46*) dataset with TALLSorts multi-class labels. Axis values are representative of the absolute number of samples in each category. **d**, Same but with TASC predictions. **e,** PCA of a patient-derived xenograft cohort (n=24), plotting PC1 and PC2 with variance explained. Points are colored by predicted subtype, with PC1 primarily reflecting subtype-driven variation. **f,** UMAP embedding of sample SCPL000710 (scRNA-seq) showing cluster assignments and coloured according to the respective cluster. **g,** UMAP from **f** annotated according to the predicted subtypes per cluster based on TASC analysis of cluster-level pseudobulk profiles. Cluster-specific colors match predicted subtype labels.

Cases harboring TAL1 or LMO2 alterations were predominantly classified into the TAL1 DP-like, TAL1 αβ-like, LMO2 γδ-like, or ETP-like subtypes, in agreement with their underlying transcriptional programs. Notably, LMO2 γδ-like cases remained detectable through probability thresholding even after imputation-associated decreases in predicted probabilities, demonstrating the robustness and practical utility of threshold-based subtype assignment (**Supplementary Data 8**). Two TAL/LMO2-altered cases were classified as SPI1, a class not originally captured by existing subtype frameworks. Subsequent gene fusion analysis revealed *STMN1::SPI1* fusions in both cases, consistent with the t (5;11) translocation and further corroborating TASC’s capacity to uncover cryptic or underrecognized subtypes (**Figure 4a**).

Overall, TASC demonstrates robust performance across datasets, achieving high subtype concordance without requiring batch correction. According to Principal Component Analysis (PCA), TASC-predicted subtypes were clearly separable along PC1 and PC2, suggesting that multi-omic subtype identity constitutes a major independent axis of transcriptional variation (**Figure 4b**).

To further validate model performance, we applied TASC to a small panel of 10 T-ALL cell lines. With the exception of CCRF-CEM, all lines were assigned to subtypes consistent with their known genomic alterations (**Supplementary Table 4**, **Supplementary Data 9**).

This provides additional support for the robustness of TASC across both primary samples as well as patient-derived and *in vitro* systems

### TASC outperforms existing tools in the identification of high-risk subtypes

To further assess the performance of TASC, we benchmarked it against the TALLSorts classifier(*45*) using two independent cohorts: one comprising 29 primary patient samples(*46*) and the other consisting of 25 patient-derived xenografts (PDXs)(*47*), both external to the training data of either classifier. In the primary sample cohort, where ETP-ALL versus non-ETP-ALL annotations were available, TASC demonstrated superior sensitivity in detecting ETP-like cases. Most ETP-ALL samples were assigned to the ETP-like subtypes by TASC, whereas TALLSorts placed only a minority of them under the “Diverse” category, despite this class being reported to cover the majority of ETP-ALL (**Fig. 4c–d**).

Although the classification of TLX1 and TLX3 subtypes remained largely concordant between the two models, major differences emerged in the assignment of TAL1-altered samples. While several were consistently classified, TASC identified additional molecularly distinct subgroups within the TAL1 spectrum, offering improved granularity (**Supplementary Table 5**). This refined separation enables TASC to more effectively delineate high-risk subtypes, such as ETP-like(*28*), which are often underrepresented or misclassified by existing methods.

Comparable findings were observed in the PDX dataset, where classification concordance between TASC and TALLSorts was generally high (**Supplementary Table 6**). Nevertheless, TASC again provided a more nuanced separation of TAL1-driven subtypes. In this cohort, principal component analysis revealed that TASC-defined subtypes aligned with the primary axis of variation (PC1), reinforcing the model’s ability to capture biologically relevant distinctions and suggesting that subtype identity is a principal driver of transcriptional heterogeneity in T-ALL (**Figure 4e**).

### Exploring intra-patient heterogeneity using TASC

To assess TASC’s ability to capture intra-patient transcriptional heterogeneity, we applied the model to a single-cell RNA sequencing (scRNA-seq) cohort comprising 41 samples, 36 of which had matched bulk RNA-seq data. The cohort was relatively homogeneous, with ETP-ALL representing 30 of 41 cases (73%). Comparison of bulk RNA-seq predictions with subtype assignments across scRNA-seq clusters showed substantial concordance: in 28 of 36 cases, the subtype predicted from bulk data matched the subtype assigned to the majority of single-cell clusters (**Supplementary Data 10**). As expected, predicted probabilities were systematically lower in the scRNA-seq setting, reflecting differences in data modality rather than loss of subtype signal (**Supplementary Figure 4a**). Evidence of intra-patient heterogeneity was observed in 14 of 41 cases, in which more than one transcriptional subtype was detected across clusters. This phenomenon was present in both ETP-ALL (**Supplementary Figure 4b–c**) and non-ETP samples (**Figure 4f–g**, **Supplementary Figure 4d–e**) and mostly involved ETP-like, TAL1, SPI1 and TME-enriched subtypes, which do not represent conflicting oncogenic programs. For example, in a TAL1 αβ-like sample, up to three distinct subtypes were identified, including both TAL1-related transcriptional programs and an ETP-like component (**Supplementary Figure 4b–c**). Interestingly, small ETP-like clusters were also detected in otherwise non-ETP samples (**Figure 4f–g**), highlighting the utility of TASC for uncovering minor transcriptional subpopulations that may represent high-risk subclones with potential relevance for disease progression or treatment resistance (48). Cell-type composition inferred using SingleR did not reveal major differences across subtype-assigned clusters, except for TAL1 DP-like and TME-enriched cases (**Supplementary Figure 4 f-h**), consistent with their known cellular composition in the original reference datasets (28). This suggests that the observed subtype heterogeneity, even with ETP-like clusters, reflects genuine transcriptional differences rather than simple shifts in cell identity.

### Leveraging TASC to unravel relapse mechanisms in T-ALL

To demonstrate the biological utility of TASC, we performed RNA sequencing on a cohort of eight matched diagnosis-relapse T-ALL samples from the Children’s Oncology Group (COG) AALL0434 clinical trial and applied TASC to obtain subtype annotations. Principal component analysis (**Figure 5a**) and surrogate variable analysis (**Supplementary Figure 5a**) revealed that subtype identity remains a major contributor to transcriptomic variance, consistent with previous analyses. Subtype predictions also highlight (in 5 out of 8 cases), the concordance between the subtype at relapse and at diagnosis (**Supplementary Table 7**) It is particularly interesting how for samples with TAL1 oncogene alterations, regardless of the transcriptional subtype, after relapse there is a drop in TAL1 class probability, which in two out of three cases is accompanied by an increase in LMO2 γδ-like subtype probability (**Figure 5a**), suggesting a switch in transcriptional programs of these non-mutually exclusive oncogenes(*26*).

**Figure 5:**
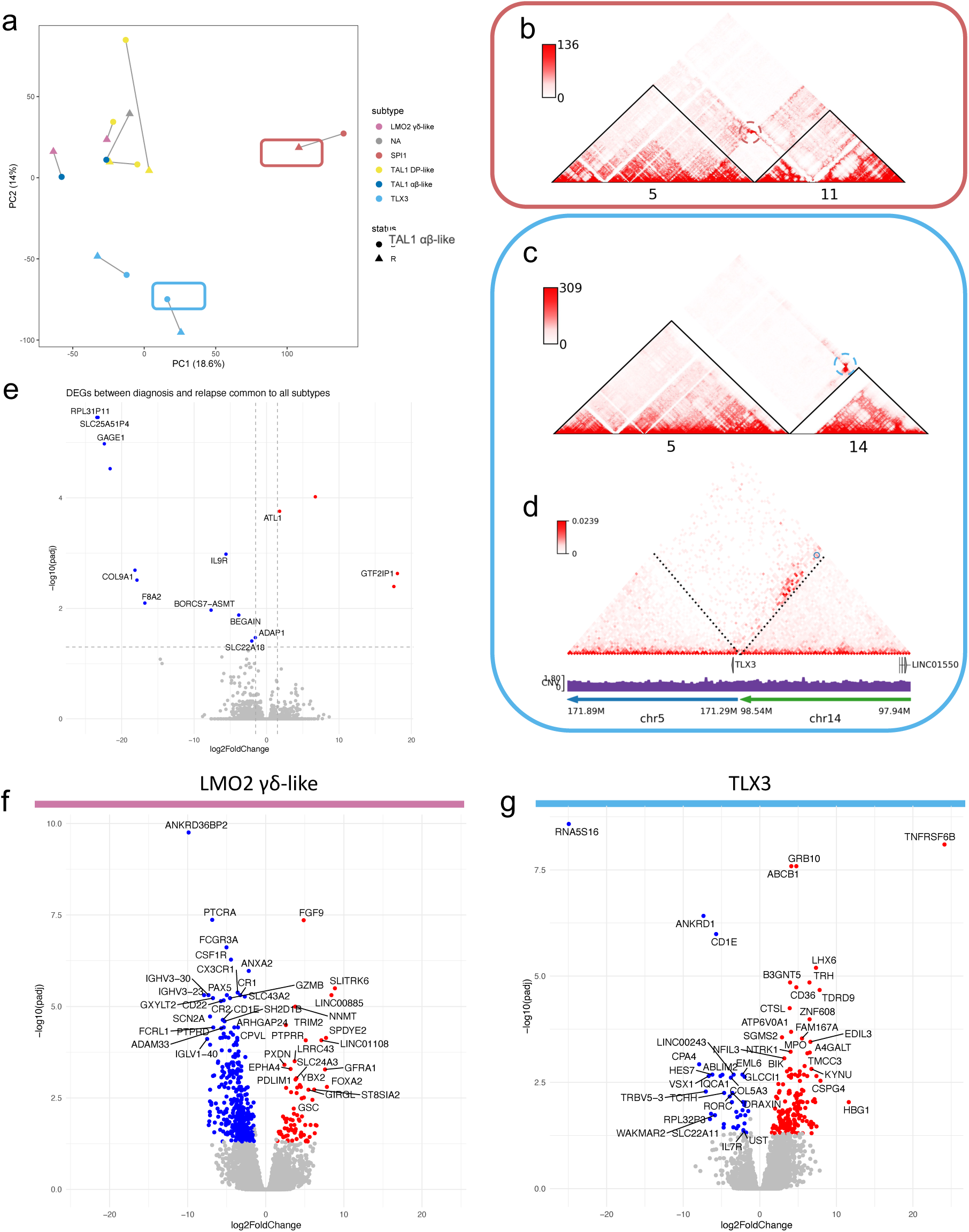
Integrative analysis of diagnosis (dx) vs. relapse (rx) samples in the dx-rx cohort unveils subtype-specific mechanisms of relapse. **a** PCA of the dx-rx cohort, showing PC1 vs. PC2 with percentage variance explained. Each point represents a sample colored by subtype. Shape is set to highlight differences between the diagnosis and relapse groups, and each pair is joined by a segment. **b,** Contact map obtained by HiChIP analysis, with colour scale bar showing distance and Copy Number Variation (CNV)-scaled interaction strength at 10 kb resolution, highlighting translocation breakpoints between chromosome 5 and chromosome 11 leading to *TCF7::SPI1* gene fusion in the relapse sample classified as SPI1. **c**, same as **b,** but for the *TLX3::LINC01550* enhancer hijacking event between chromosome 5 and chromosome 14 observed in one of the diagnosis primary-samples classified under TLX3 subgroup. **d**, Contact map obtained by HiChIP analysis, with colour scale bar showing distance and CNV-scaled interaction strength at 10 kb resolution mapped onto patient-specific reconstructed linear genome, highlighting the enhancer hijacking event in **c**, annotated with hg38 gene annotations. **e**, DESeq2 derived volcano plot representing differentially expressed genes across diagnosis and relapse groups common to all subtypes. x-axis: log₂ (fold change); Y-axis: –log₁₀ (adjusted p-value). Genes meeting significance thresholds (|log₂FC| > 1.5; –log₁₀ (adj p) > 1.3) are labeled using the corresponding gene name. Upregulated genes in relapse are shown in red; downregulated in blue. Thresholds are marked by dashed lines. **f**, Similar to **e**, but representing differentially expressed genes across diagnosis and relapse specific for the TAL1 samples switching toward LMO2 γδ-like subtype in the merged cohorts. **g**, Similar to **f** but representing differentially expressed genes across diagnosis and relapse specific for the TLX3 subtype in the merged cohorts.

To validate these findings, we performed H3K27ac HiChIP on two patients and, by applying EagleC(*49*) confirmed that the predicted subtypes were supported by structural variants identified from chromatin capture data. Notably, in one patient, the SPI1 subtype prediction was further supported by the detection of a t (5;11) rearrangement, resulting in a *TCF7::SPI1* fusion (**Figure 5b**, **Supplementary Figure 5b**). In a TLX3 subtype patient, although a t (5;14) rearrangement did not directly involve the *TLX3* locus (**Figure 5c**), HiChIP interaction maps revealed a neo-loop connecting the *TLX3* promoter with the distal non-coding element *LINC01550*, consistent with enhancer hijacking(*50*) (**Figure 5d**).

While differential gene expression analysis using subtype as a covariate, to detect shared mechanisms of relapse across subtypes, yielded little significance (**Figure 5e**), subtype-specific analysis of relapse uncovered more extensive and biologically meaningful transcriptional changes: TAL1 DP-like (n = 45 DEGs), TAL1 αβ-like (n = 399), and TLX3 (n = 82) (**Supplementary Data 11**).

However, as subtype aware analysis strongly reduces the number of samples per group, we performed integration with external paired diagnosis-relapse cohort GSE160298 after batch correction and subtype prediction. PCA after batch correction on the merged vst counts highlights the high similarity of samples with identical subtypes across the two data sets (**Supplementary figure 5c**).

Interestingly, subtype switching at relapse was also observed in this independent dataset, affecting two samples initially classified as TAL1 subtypes (one TAL1 αβ-like and one TAL1 DP-like), further supporting the biological relevance of this phenomenon. To investigate the molecular basis of subtype switching, we compared relapse-associated transcriptional changes across switching cases from both datasets. This analysis revealed a predominant downregulation of gene expression accompanied by strong relapse-associated transcriptional signatures (**Figure 5f**, **Supplementary Figure 5d**). Notably, genes involved in MHC expression and antigen presentation were significantly enriched among the downregulated genes at relapse, together with reduced JAK/STAT signaling downstream of IL-6 and IL-2, suggesting an overall attenuation of inflammatory and immune-related programs.

In contrast, relapse samples classified within the TLX3 subtype exhibited upregulation of IL-2/STAT5 signaling, enrichment of PRC2 target genes, and activation of hematopoietic stem cell–associated transcriptional programs (**Figure 5g**, **Supplementary Figure 5e**), consistent with chromatin remodeling and reactivation of stemness-related pathways (51).

Together, these results indicate that TASC not only robustly assigns transcriptional subtypes at relapse but also enables the identification of distinct, subtype-specific relapse trajectories, providing insight into the biological programs underlying disease evolution.

## Discussion

T-ALL is a genomically complex malignancy with emerging molecular subtypes, but a single unified classification framework is lacking. Therefore, we developed TASC, an RF-based classifier that uses RNA sequencing data to recognize established and clinically relevant subtypes in T-ALL cases. TASC provides a unified transcriptional classifier for up to 13 T-ALL subtypes. Trained on large multi-cohort data on widely accepted multi-omic subtypes (*28*), it performs class prediction with high global accuracy (0.92) based on a set of 300 highly variable genes across subtypes. In contrast to existing tools that often rely on linear models such as one-vs-rest logistic regression, TASC employs a RF ensemble, enabling the capture of non-linear gene–gene interactions and improving robustness to noise and class imbalance, similar to other approaches used in B-ALL (*53*). Compared to classifiers based on different feature sets and more complex gradient-boosted models, TASC demonstrates superior performance, illustrating that model simplicity, when combined with informative features, can effectively address multi-class subtype prediction.

TASC provides interpretable predictions through both global and local feature importance analyses(*54*), consistently highlighting subtype-specific transcription factors and oncogenes(*5*), alongside additional genes implicated in T-ALL pathogenesis. For example, genes such as *HPGDS* and *MN1*, associated with metabolic regulation and stemness(*41–44*), emerged as putative predictors of the ETP-like subtype. These findings enhance our understanding of the less characterized subgroups and further underscore the biological relevance of TASC’s decision-making framework.

While classification performance varies across subtypes – particularly in low-incidence groups excluded from the final classifier, such as NKX2-5 and STAG2/LMO2 - this variation reflects both the class imbalance present in current datasets and the biological ambiguity of certain transcriptomic profiles. Nonetheless, TASC effectively distinguishes most rare cases and flags low-confidence predictions with lower posterior probabilities, which facilitates downstream interpretation. This suggests that future efforts to improve representation of rare subtypes (e.g. by augmenting sample sizes through additional patient data collection) could significantly enhance TASC sensitivity and stability for low-incidence classes. Furthermore, the framework’s simplicity and flexibility allow for easy retraining as larger, more curated datasets become available or new subtypes are reported. This addresses a key limitation of current methods and offers a scalable approach for continuous refinement. TASC was benchmarked against the only available T-ALL transcriptome-based classifier(*45*) using independent datasets comprising both primary patient samples and patient-derived xenografts. Across these external cohorts, TASC demonstrated high overall concordance with published subtype annotations. Importantly, TASC showed increased sensitivity for high-risk subtypes, particularly for ETP-ALL, and was able to improve the stratification of cases from previously merged categories(*45*), such as samples displaying *TAL1, LMO2, KMT2A, MLLT10*, and *HOXA9* abnormalities. Beyond primary samples, TASC also performed well on additional sample types, including RNA-seq from T-ALL cell lines and PDX samples, proving accurate predictions across diverse datasets and sample types. By allowing subtype prediction of high-risk subtypes, TASC can streamline and enhance diagnostic workflows and translational research in T-cell acute lymphoblastic leukemia(*21*). Furthermore, TASC can be applied to single-cell RNA-seq data to resolve intra-tumoral heterogeneity and infer subtype composition at cellular resolution. Finally, in matched diagnosis–relapse cases, TASC captures subtype conservation and highlights relapse-associated transcriptional dynamics, supporting its potential utility for studying disease evolution and therapeutic resistance. Given the limited number of available paired samples (four subtype-switching and four TLX3 pairs), these findings primarily serve as proof-of-concept and warrant validation in larger longitudinal cohorts(55).

The classifier is made accessible through a user-friendly Shiny web application, available both online and for local deployment on institutional servers. This flexibility lowers the barrier to adoption, allowing researchers and clinicians to obtain subtype predictions in a straightforward and reproducible manner. Our tool accepts inputs in the form of raw gene expression counts generated by commonly used RNA-seq processing pipelines, including featureCounts(*56*), Salmon(*57*), and nf-core/rnaseq(*58*), facilitating its seamless integration into existing bioinformatics workflows. In addition to subtype predictions and associated posterior probabilities, TASC provides interactive visualizations, including principal component analysis and metagene plots based on differentially expressed genes across subtypes, to evaluate the concordance of the predictions with the known gene expression profiles across classes. To ensure robustness, the tool includes automated imputation for up to 10% of missing predictor genes, reducing the risk of unreliable classification due to incomplete data while maintaining predictive accuracy. In addition to maximum-probability subtype assignments, TASC provides relative probability tables and supports filtering based on clinically informed probability thresholds, enabling more nuanced interpretation of classification confidence.

An important limitation of this study is the strong class imbalance inherent to T-ALL transcriptional subtypes, which reflects true biological rarity rather than insufficient cohort size. Several subtypes are represented by fewer than ten samples, precluding reliable stratified cross-validation without introducing artificial resampling or data leakage. For this reason, we prioritized external validation across multiple independent datasets rather than cross-validation within the training set, as repeated resampling of extremely rare subtypes would be expected to inflate performance estimates and reduce biological interpretability.

Additional limitations should be noted. A further limitation is that TASC was trained exclusively on bulk RNA-seq data and was not optimized for single-cell transcriptomic inputs. As a result, subtype predictions at the single-cell level exhibit reduced probability confidence and lower concordance with bulk assignments, reflecting both technical sparsity and biological heterogeneity inherent to scRNA-seq data. Consequently, TASC does not directly identify genetic subclones but instead highlights transcriptionally distinct cellular populations whose expression profiles are consistent with known subtype programs. While this application enables the exploration of intra-patient heterogeneity and the identification of minor subtype-like populations, definitive subclonal inference will require integration with single-cell genomic or multi-omics approaches in future studies.

Finally, although TASC was validated across multiple independent datasets, the availability of large external cohorts with complete transcriptomic profiles that do not require gene imputation remains limited. As a result, validation of some datasets relied on partial imputation, which—while systematically evaluated—may attenuate prediction confidence. Future studies leveraging larger, prospectively collected cohorts with comprehensive transcriptomic coverage and minimal missing data will be essential to further strengthen validation and expand clinical applicability.

By providing comprehensive subtype classification from a single RNA-seq assay, TASC has the potential to replace multiple targeted tests, paralleling recent advances in the B-ALL field, where transcriptome-based classifiers are increasingly becoming standard practice (*5*, *22*, *23*). However, its deployment in routine diagnostics may require simplification: although RNA-seq offers powerful insights, a focused gene panel based on TASC’s key predictors could be developed for more practical use. Future work will address these challenges by incorporating larger, clinically-annotated cohorts (including relapse cases) and prospectively evaluating TASC’s prognostic value. These efforts will help refine TASC into a robust and clinically practical framework for T-ALL subtype classification.

## Methods

### Public cohorts and dataset

This work includes analysis of public RNA-seq and scRNA-seq datasets, comprising a heterogeneous set of samples, including primary samples, cell lines, and PDX. Source, references, and downloaded data type are reported (Supplementary Data 1).

### Paired diagnosis-relapse cohort

Human blood samples used for RNA-seq, H3K27ac HiChIP analyses were obtained from the Children’s Oncology Group (COG) AALLNV05_Q protocol and approved by the COG Disease Biology Subcommittee. Bone marrow aspirates were collected and immediately processed to select mononuclear cells by Ficoll-Paque PLUS density gradient medium (GE Healthcare, 17-1440-02) per the manufacturer’s instructions. Samples were then either used fresh or vitally frozen in 10% DMSO, 90% serum for subsequent use.

### RNA sequencing

RNA was extracted from sorted samples using RNEasy plus micro kit (Qiagen). To allow for the detection of mRNAs and ncRNAs from low-input samples, RNA-seq libraries were prepared using NuGEN Ovation Trio (Tecan) following the manufacturer’s guidelines. Libraries were sequenced in paired-end by NovaSeq-6000 on SP flow-cell at 300 cycles. Raw RNA sequencing data were processed using the nf-core/rnaseq pipeline (v3.14.0) with the Singularity execution profile. The workflow was run with STAR for alignment and Salmon for quantification (--aligner star_salmon), using the *Homo sapiens* GRCh38 reference genome (Ensembl release 112). Custom STAR alignment parameters were applied to adjust read filtering thresholds (--outFilterScoreMinOverLread 0.3, --outFilterMatchNminOverLread 0.3). The pipeline was executed with gene annotation from GENCODE and duplicate rate estimation (dupradar) disabled.

### Sequencing of H3K27ac HiChIP

HiChIP was performed using the Arima HiC+ kit (Arima Genomics) following manufacturer’s instructions (March 2020 user guide), with some modifications. Briefly, up to 3 million cells were crosslinked in 2% Formaldehyde (Sigma-Aldrich) for 10 minutes at room temperature. Formaldheyde was neutralized by adding Stop Solution 1 (Arima HiC+ kit) and processed following manufacturer’s instructions for steps: (1) restriction enzyme digestion, (2) chromatin shearing, (3) antibody incubation, (4) protein A beads blocking, (5) IP and washes (6) quality controls, (7) library preparation, (8) library complexity QC and (9) library amplification. For step 2, chromatin was sheared on Covaris ME 220 (Covaris) at 100W peak incident power, 10% duty factor, 200 cycles per burst, 300sec treatment time, 4 degrees Celsius. For step 5, samples were immunoprecipitated using Abflex Histone H3K27ac antibody (#91193 ActiveMotif, RRID: AB_2793797) in ratios calculated from the Arima Sample Sheet. Libraries were sequenced in paired-end by NovaSeq on SP flow-cells at 100 cycles per read.

## Statistical analysis

### Surrogate variable analysis

To identify and account for latent sources of variability not attributable to the condition of interest (diagnosis vs. relapse), surrogate variable analysis was performed using the DaMiRseq R package (v1.12.0), following the recommended pipeline. Variance-stabilized gene expression data were used as input, and surrogate variables (SVs) were inferred using partial least squares regression. To investigate the nature and potential collinearity of these SVs with known biological or technical factors, Spearman correlation coefficients were calculated between each SV and sample-level covariates, including condition (diagnosis vs. relapse), predicted T-ALL subtype, and patient. Results were visualized as a correlogram.

### Differential expression analysis and functional enrichment

Differential gene expression analysis in the Diagnosis:Relapse cohort was performed on *salmon* gene-level quantifications from eight paired diagnosis–relapse samples, integrating subtype prediction from the TASC web-based Shiny app. Relapse-associated, subtype-independent transcriptional changes were assessed using the DESeq2 R package (v1.44.0), employing a no-intercept model with subtype included as a covariate. In parallel, subtype-specific changes were assessed by merging our internal dataset with external paired diagnosis-relapse cohort GSE160298 (where TASC was used to determine subtypes on the raw-counts, after imputation of 25 genes), performing subtype agnostic batch correction with R package CombatSeq. The combined dataset, annotated according to predicted subtypes, was used for differential expression analysis with DESeq2 under a no-intercept model. Subtypes with fewer than two pairs were excluded from subtype-specific analyses, and class-switching samples were analyzed separately. Significant genes (|log₂ fold change| > 1.5, adjusted *p* < 0.01) were categorized into upregulated and downregulated sets and used for functional enrichment analysis with the EnrichR R package (v3.4). Enrichment was assessed across the following gene set libraries: MSigDB Hallmark 2020, GO Molecular Function 2023, GO Cellular Component 2023, GO Biological Process 2023,

KEGG 2021 Human, BioCarta 2015, Panther 2015, WikiPathways 2024, Reactome 2024, ENCODE TF ChIP-seq 2015, ChEA 2022, and CellMarker 2024. Enriched terms were considered significant if they met the thresholds of adjusted *p* < 0.05, gene set intersection size > 3, and odds ratio > 1.

### Data analysis from external cohorts

Fastq files from the CUTTL1 and CUTTL3 cell lines, and primary samples GSM3004557 and GSM3004625 were aligned to the GRCh38.p14 genome using custom scripts and parameters (https://github.com/CBenetti/RNAVardetect), using STAR (2.7.11). We then marked duplicate reads using *Picard* (2.26.10) *MarkDuplicates*, tagging PCR and optical duplicates based on alignment positions to prevent bias. Read groups were assigned with *Picard AddOrReplaceReadGroup***s**. Spliced RNA alignments were processed using *GATK* (4.2.1.0) *SplitNCigarReads*, which splits reads at splice junctions and adjusts CIGAR strings to correct for intron-spanning alignments. Base quality scores were recalibrated through *GATK BaseRecalibrator* using known variant databases, followed by *GATK ApplyBQSR* to correct systematic sequencing biases. Raw counts for T-ALL cell lines CUTTL1 and CUTTL3 were then obtained using featureCounts (subread, 1.6.3) using Ensembl GRCh38.p14 reference annotations and merged with other cell line datasets. Variance stabilizing transformed (vst) counts were obtained through DESeq2 (v1.44.0).

### Gene fusion detection from RNA sequencing

Starting from the processed Chimeric junction file from the RNA-seq data analysis pipeline, star-fusion (1.9.0) was used to detect gene-fusion events on GSM3004557 and GSM3004625 using default parameters and GRCh38_gencode_v37 genome library. Visualization and plots of gene fusion events were obtained through the Chimeraviz R package (1.24.0) using Ensembl Homo_sapiens.GRCh38.112 transcripts as references

### Computational analysis of HiChIP data

H3K27ac HiChIP sequencing data from T-ALL cell lines were processed using the MAPS-Arima pipeline. Reads were aligned to the hg38 reference genome and filtered via the feather module. Peaks were identified using MACS2 with MAPS-specific parameters optimized for HiChIP data. Pairtools was employed to generate paired-end interaction files to enable downstream chromatin interaction analysis, which were subsequently used as input for the HiC-bench framework. Contact matrices were generated with Juicer and converted to the “.cool” format using hic2cool. To correct for distance-dependent interaction biases and copy number variation (CNV), matrices were normalized using NeoLoopFinder at 10 kb resolution

### Structural variant identification

Distance-CNV normalized CNV profiles were used to detect structural variants from HiChIP contact maps. Prediction was performed using EagleC, and the results were used as input to NeoLoopFinder to detect patient-specific genomic assembly and the related neoloops. Chromatin interaction maps were generated using EagleC and NeoloopFinder plotting utilities.

### Analysis of single cell RNA sequencing

Raw scRNA-seq data from each sample’s *SummarizedExperiment* were converted as Seurat objects Data were centered and scaled using Seurat’s *ScaleData ()* function on previously identified variable features (2,000 genes). Features were centered at zero mean and scaled to unit variance. Extreme values were clipped with a maximum absolute value of 10 to reduce the influence of outliers. No variables were regressed out. A shared nearest neighbor graph was constructed using *FindNeighbors ()* with principal components 1–10 and default settings: k.param = 20, Jaccard-based SNN, prune.SNN = 1/15, and exact nearest neighbor search (nn.method = “rann”). Communities were then identified using *FindClusters ()* with the following parameters: resolution = 0.3, algorithm = 1 (standard Louvain), modularity function = 1, n.start = 10, n.iter = 10, and singleton cell grouping enabled. Cluster cell composition was defined by applying SingleR to assign labels for each single cell according to BluePrint and HumanCellAtlas merged annotations, and then aggregated based on clusters. Cluster-specific pseudobulk profiles were generated by aggregating single-cell data per cluster. Pseudobulk profiles were subsequently classified using TASC, providing insights into subtype-specific heterogeneity at the cellular level. Predicted probabilities were compared for clusters whose subtype assignments agreed with the corresponding bulk RNA-seq data.

### Selection of model predictors

Two sets of marker genes were used as model features. The 300 most variable genes (MVGs) used for subtype clustering in the Pölönen et.al. publication(*28*), confirming substantial overlap with the top 300 variable genes in the training set, calculated by standard deviation on transcript per million (TPM) batch corrected and filtered counts (excluding genes with expression of more than 1 TPM < 5 samples). Subtype-grouped dendrograms were generated using variance-stabilized transformed (vst) expression profiles of MVGs. Euclidean distance and complete linkage clustering were applied to these expression profiles to evaluate subtype relationships.

Alternatively, subtype-specific markers were calculated similarly to what was previously suggested in Pölönen et.al. publication, by performing Differential Gene Expression (DGE) analysis for each pairwise permutation of subtypes using EdgeR (4.2.2). Counts were normalized using the trimmed mean of M-values (TMM) method implemented in *edgeR*’s *calcNormFactors ()* function. Quasi-likelihood testing with *glmQLFit ()* without an intercept was used to assess pairwise subtype comparisons. Significant results (*FDR < 0.01, abs (log fold change) >1*) were assigned to either of the compared groups based on logFoldChange sign and mean expression of 2 TMM in the upregulated subset. After merging results from different comparisons of the same class, unique genes were selected as subtype markers.

### Random Forest model tuning

The original cohort was randomly split into training (n=850, 64%) and testing (n=475, 36%) sets. Samples belonging to subtypes with very low representation (<1% frequency) were excluded from each set independently. To evaluate classification performance, a custom summary function was defined to calculate sensitivity and specificity using the first factor level as the positive class. This function was integrated into the *trainControl ()* function from the caret (7.0-1) package using 10-fold cross-validation with class probability tracking enabled. Random Forest models were trained using the randomForest (R, “4.7-1.2”) function on variance-stabilized normalized data, using defined sets of predictor features and the following hyperparameters: ntree=500, nodesize = 1 and nodesize = *sqrt(p)*, where *p* is the number of predictors for each set of selected features. After an initial training round, weights for underperforming subtypes (sensitivity <0.9) were increased to 10 to improve recall, followed by retraining using weights and importance = TRUE. Classification performance, including F1-score, precision, recall, and balanced accuracy, was assessed using the *confusionMatrix ()* function, comparing true labels with predicted outputs in both training and testing sets. Probabilities for the positive class were extracted from the trained random forest in the testing set, and PR-AUC values were calculated using the PRROC package (1.4) *pr.curve ()* function for PR analysis while p-AUCs and probability thresholds for the MVG model were computed using *roc()* function from *pROC* package in R. For each sample, the predicted class probabilities of the test set were used to determine the confidence (maximum predicted probability) and the predicted label (class with maximum probability). Predictions were grouped into 5 equally spaced confidence bins between 0 and 1. For each bin, the average confidence, observed accuracy, and 95% confidence intervals (via Wilson method) were calculated. The ECE was computed as the weighted average of the absolute differences between average confidence and accuracy across bins. Model calibration was further assessed using the Brier score, which was computed separately for the training and test sets and for each model, as the mean of the row-wise squared differences between predicted probabilities and true labels.

### Comparative model evaluation

We trained a multiclass gradient-boosted tree model using the XGBoost R package (1.7.11.1). Key hyperparameters were set as: learning rate of 0.01, maximum tree depth of 8, gamma value of 4, subsampling rate of 0.75, and full column sampling. The objective function was set to output class probabilities (multi:softprob), evaluated using multiclass log-loss. The model was trained for 5,000 boosting rounds on variance-stabilized gene expression data formatted as an xgb.DMatrix. Model input features, training, and testing set composition were set to match the ones used for TASC random forest classifier. Both models were evaluated on the same held-out test data using comprehensive performance metrics, including F₁-score, precision, recall, and balanced accuracy. Metrics were derived via *the confusionMatrix ()* function from the *randomForest* package.

### Validation and benchmarking

Performance was rigorously assessed on the independent testing dataset and further validated against multiple external RNA-seq datasets, including 499 B-ALL subtype-annotated samples, used as negative-validation, 80 primary samples with detailed genomic annotations from GEO datasets GSE110633 and GSE110636; 10 T-ALL commercial cell lines from GEO dataset GSE103046, in addition to CUTTL3 (GSE243914) and CUTTL1 (GSE59810) cell lines; 24 primary samples with ETP status annotations from GEO dataset GSE243914. Benchmarking with existing RNA-seq classifier TALLSorts (https://github.com/Oshlack/TALLSorts.git) was performed by comparing predicted annotations from TASC to TALLSorts predictions on datasets external to both algorithms, specifically GEO dataset GSE243914 and a dataset of 25 PDX samples. In the entirety of these cases, TASC predictions were obtained from web-based Shiny App, including an imputation step for missing features (https://01971d1c-d81d-a598-4238-5afb7b3e381a.share.connect.posit.cloud/).

### AI explainability methods

Global feature importance was assessed by calculating feature cross-entropy loss using the iml R package (0.11.4). Local explainability was assessed using SHAP values. For each class, a custom prediction wrapper was defined to extract class-specific probabilities. The SHAP values were computed with the fastshap package (0.1.1), leveraging parallel computation to enhance efficiency. The baseline prediction was set as the average predicted probability for the class across the training data. Resulting SHAP values quantified the impact of each gene on the model’s output, enabling a detailed interpretation of feature importance at the sample level. Dendrograms were constructed from the median of scaled SHAP values to reflect model-driven subtype similarities. For these, Pearson correlation distance and Ward’s D2 linkage method were used. Global SHAP contribution was calculated by averaging and ranking local values for each class, and spearman correlation was performed to determine the concordance with mean decrease in Gini coefficient.

### Evaluation on batch and imputation contributions

Gene expression data from the testing set, datasets GSE110633 and GSE110636 (external validation cohort) and bulk RNAseq data from the matched scRNA-seq cohort were preprocessed and variance-stabilized using DESeq2.

On the test set, a 30 feature sliding-window permutation and incremental random feature imputation by sampling from normal distributions parameterized by the mean and standard deviation of the training set were performed to asses the effect on prediction probabilities.

Batch effects across datasets were assessed using principal component analysis (PCA) on combined expression matrices. PCA plots were annotated with sample subtype and batch information, and centroids per batch were computed. For the external validation cohort, missing values in selected gene subsets were imputed by sampling from normal distributions parameterized by the mean and standard deviation of testing and training data genes. To quantify batch contributions (in both cohorts, with and without imputation) to prediction probabilities, linear models were fit for each subtype with predicted probabilities as the dependent variable and the dataset as the independent variable. Subtypes represented in more than one dataset were included, and non-intercept coefficients were extracted to evaluate systematic batch effects dependent on batch alone or batch and imputation (GSE110633 and GSE110636).

## Acknowledgements

We acknowledge all members of the Boccalatte laboratory for helpful discussions. We would like to thank Guido Barzaghi and Giorgio Inghirami for their critical reading of the manuscript. We thank the Genome Technology Center (GTC) for library preparation and sequencing, partially supported by the NYU Cancer Center Support grant P30CA016087 at the Perlmutter Cancer Center. This work has used computing resources at the NYU Grossman School of Medicine High Performance Computing (HPC) Facility. We also thank the Candiolo Genomics Center for library sequencing and troubleshooting. We would also like to thank Hansen Kosasih and Charles De Bock (UNSW Sidney) for sharing raw datasets for xenografts.

## Author contributions

Conceptualization: CB, FB

Methodology: CB, OA, GG, AM, FB

Investigation: CB, OA, GG, AM, FB

Data curation: IA, AT, FB

Supervision: IA, AT, FB

Writing—original draft: CB, FB

Writing—review & editing: CB, FB, OA, GG, AM, IA, AT

## Data and code availability

The publicly available NGS datasets used for the present study are listed in Supplementary Data 1. All sequencing data created in this study (RNA-seq and HiChIP) will be deposited in the Gene Expression Omnibus (GEO; https://www.ncbi.nlm.nih.gov/geo/) repository [accession code pending]. All the code from this project uses published packages. TASC source script is available on GitHub in the following repository: https://github.com/CBenetti/TASC. TASC is also available as a web-hosted Shiny App in the following repository: https://01971d1c-d81d-a598-4238-5afb7b3e381a.share.connect.posit.cloud/. Details of the modules added to the original <u>NYU-BFX/**hic-bench**</u> framework for alignment and structural variant analysis on HiChIP data can be found at the following link: https://github.com/CBenetti/hic-bench-arima.

