## Supplementary Figures and Tables for "TASC: A transcriptome-driven machine learning classifier to explore molecular heterogeneity and relapse-associated programs in T-cell Acute Lymphoblastic Leukemia"

**This PDF file includes:**

Supplementary Figures. SF1 to SF5
Supplementary Tables ST1 to ST7

**Other Supplementary Materials for this manuscript include the following:**

Supplementary Data SD1 to SD11

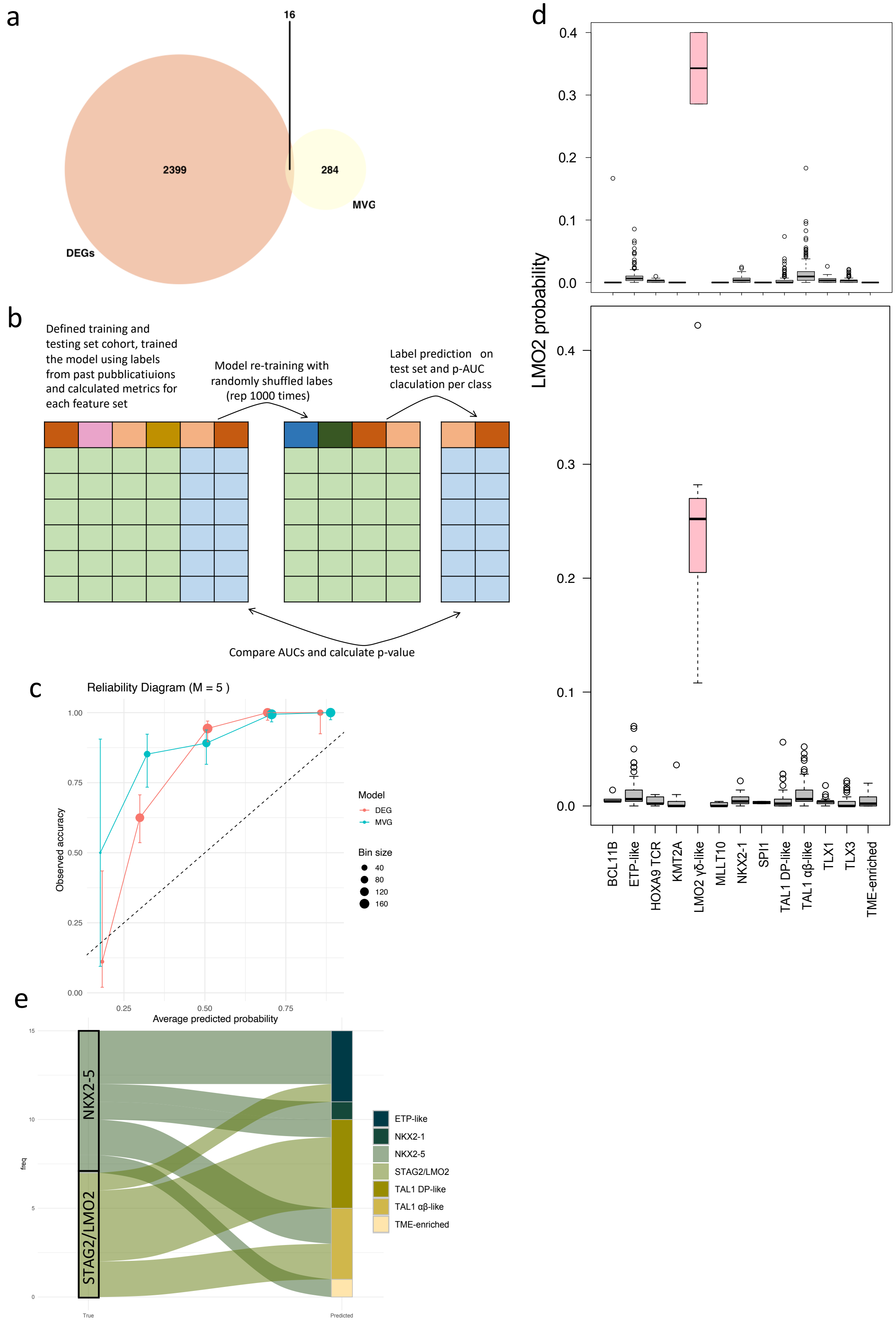
a
16
2399
284
MVG
DEGs
b
Defined training and testing set cohort, trained the model using labels from past pubblicatiuions and calculated metrics for each feature set
Model re-training with randomly shuffled labes (rep 1000 times)
Label prediction on test set and p-AUC claculation per class
Compare AUCs and calculate p-value
c
Reliability Diagram (M = 5 )
Observed accuracy
Average predicted probability
Model
DEG
MVG
Bin size
40
80
120
160
d
LMO2 probability
BCL11B
ETP-like
HOXA9 TCR
KMT2A
LMO2 γδ-like
MLLT10
NKX2-1
SPI1
TAL1 DP-like
TAL1 αβ-like
TLX1
TLX3
TME-enriched
e
NKX2-5
STAG2/LMO2
freq
True
Predicted
ETP-like
NKX2-1
NKX2-5
STAG2/LMO2
TAL1 DP-like
TAL1 αβ-like
TME-enriched

**TASC calibration and metrics**. **a**, Venn diagram of intersection between the two set of predictors: differentially expressed genes (DEGs, pink) and the 300 most variable genes (MVG, yellow). **b,** Schematic representation of workflow for label permutation step used to validate model calibration. **c,** Scatterplot representing model calibration for DEG (pink) and MVG (blue) models. On the x axis the average classification probability and on the y axis the average accuracy for the 5 bins defined in expected calibration error calculation. **d**, Boxplot of LMO2 γδ-like subtype probability calculated by the MVG-based Random Forest (RF) model, split according to the true class labels of both training (green) and testing (blue) sets. Probability box from LMO2 γδ-like samples is pink-coloured, to highlight the difference in the true-class probability distribution. **e**, Alluvial plot outlining associations between original class ("True" strata) and predicted subtypes ("Predicted" strata) for samples belonging to excluded subtypes. Colors in the plot are representative of subtypes, both original (flows) and predicted. Frequency of associations is displayed on the y-axis.

**Supplementary Figure 2.**

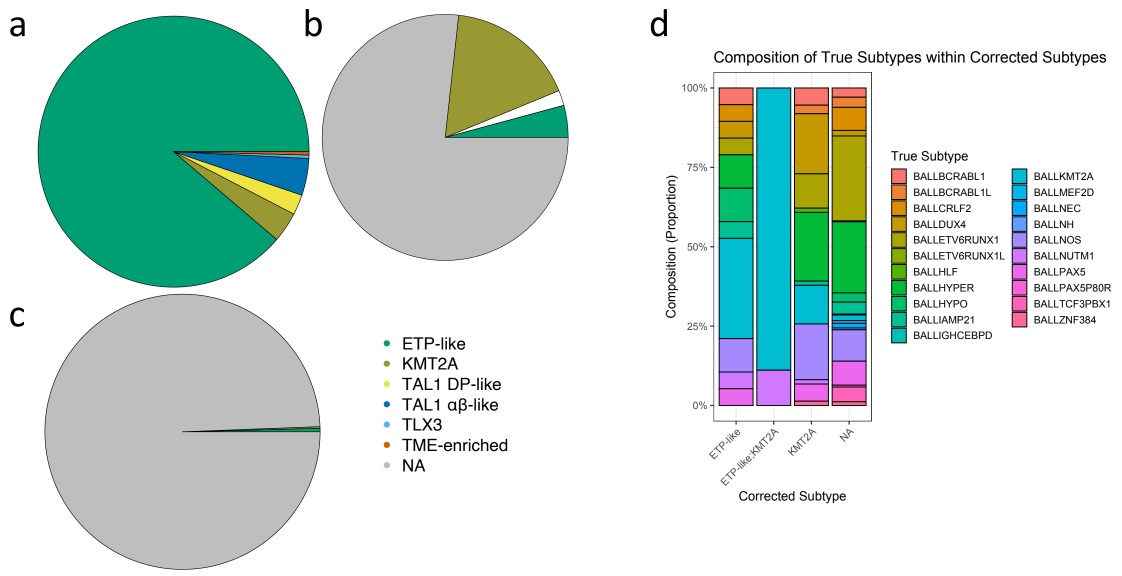


**Model tuning and validation on negative-control cohort of B-ALL patients. a-c,** Pie charts representing TASC subtype predictions on B-ALL cohort based on maximum probability calls (**a**), Youden's maxima threshold applied to probabilities (**b**) and clinically oriented custom threshold maximizing specificity (**c**). **d**, B-ALL ground truth subtype composition across predicted labels using Youden's maxima threshold on classification probabilities.

#### Supplementary Figure 3

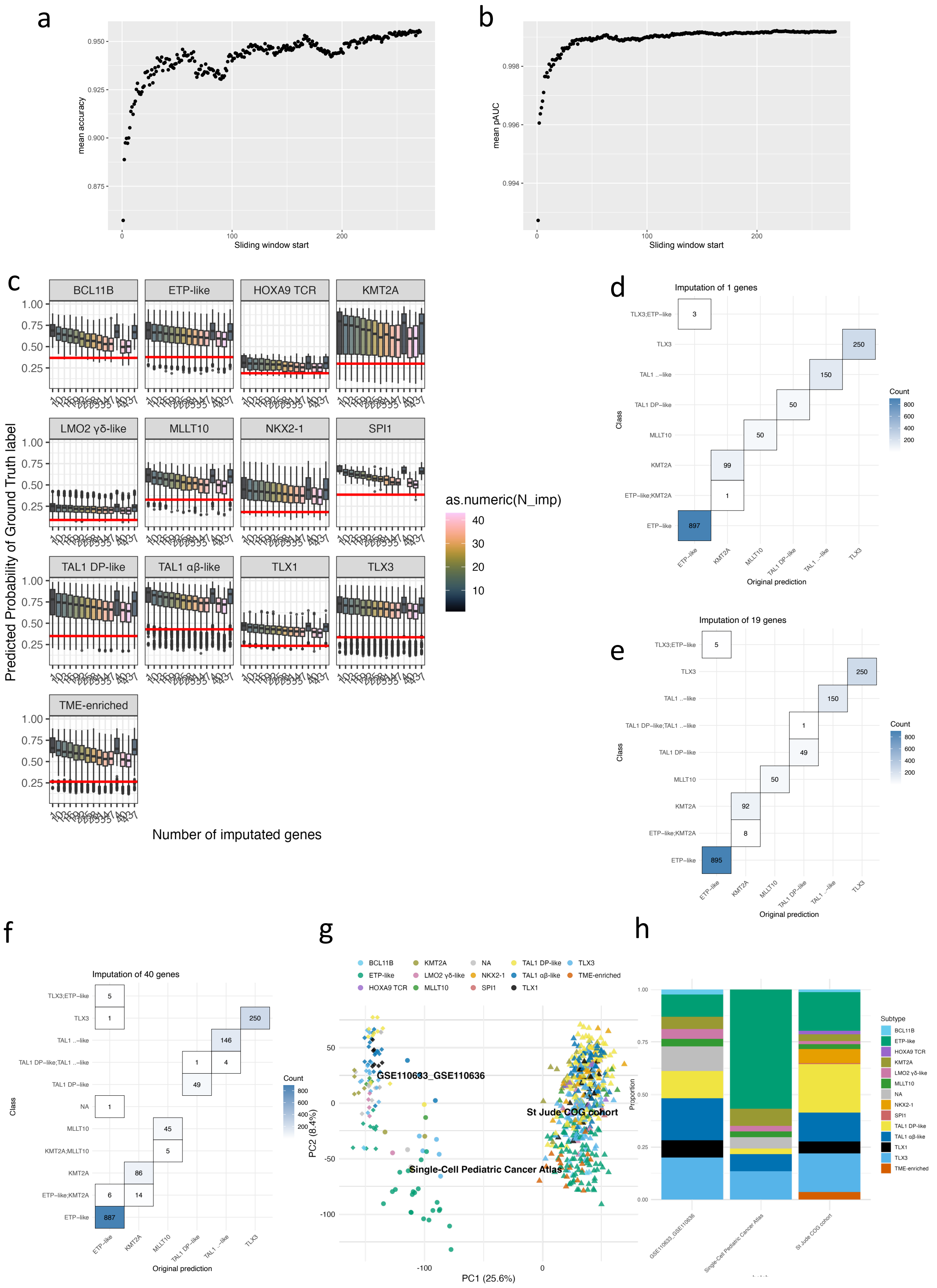

**Feature perturbation effect on accuracy and prediction probabilities. a,** Scatter plot of mean accuracy after feature permutation in a 30-gene wide sliding window in the testing set**. b,** Scatter plot of mean probability AUC after feature permutation in a 30-gene wide sliding window in the testing set. **c,** Boxplot of test-set predicted probabilities after progressive imputation of random features based on a normal distribution calibrated on the mean and standard deviation of the train set, split by subtype. **d,** Confusion matrix comparing subtype predictions without feature imputation with predictions after imputation of 1 gene based on a normal distribution calibrated on the mean and standard deviation of the train and test set **. e,** same as **d** but comparing predictions after imputation of 19 features**. f,** same as **e** but comparing predictions after imputation of 40 features. **g,** PCA plot of test set, external validation cohort and scRNAseq matched bulk RNA-seq cohort highlighting batch effect among datasets. Color is determined by the predicted subtype, while different shapes reflect the different dataset of origin**. h,** Bar plot of dataset composition, comparing predicted subtype proportions of the cohorts in **g.**

### Supplementary Figure 4

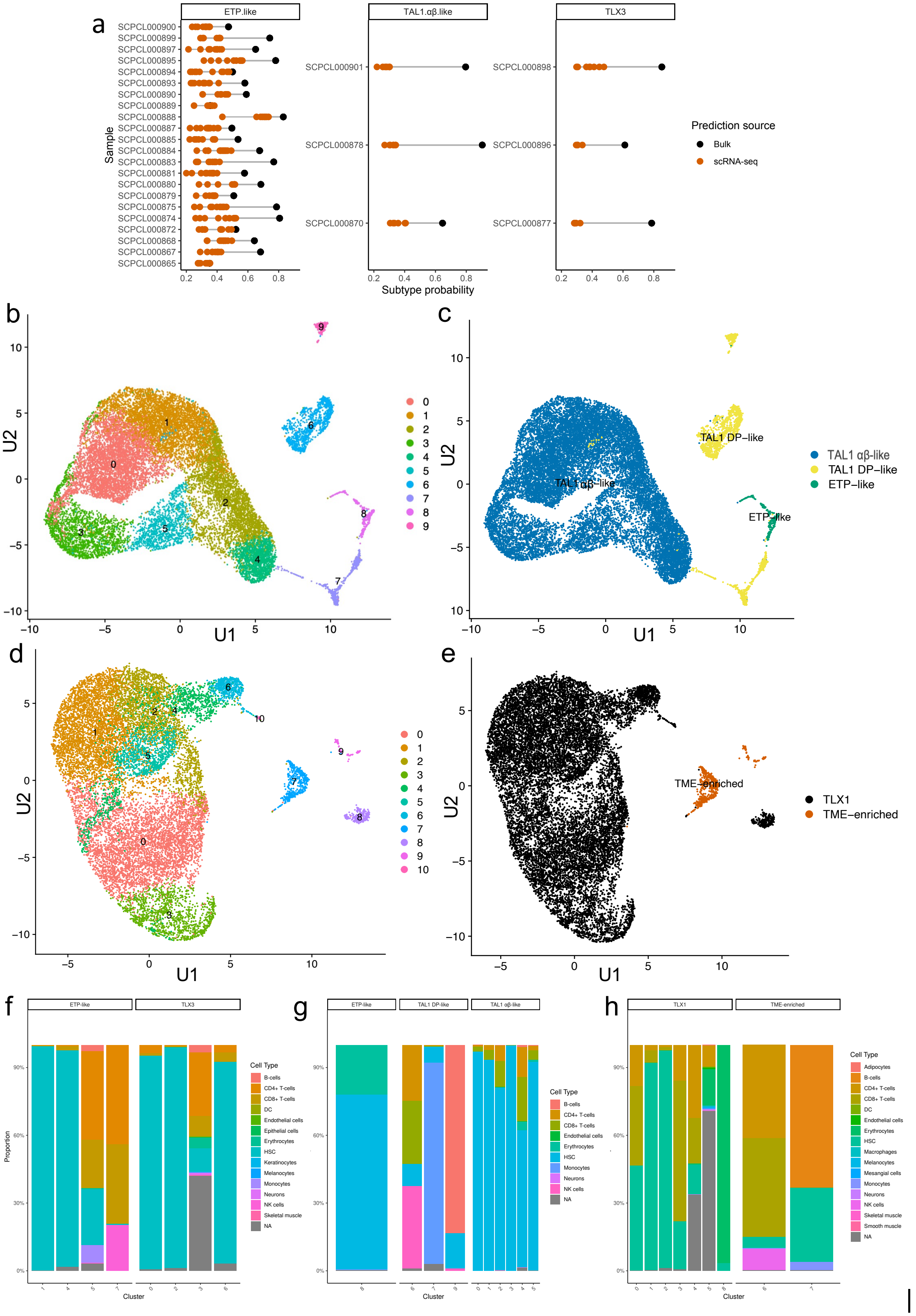

**Single-cell RNA-seq reveals intra-patient transcriptional diversity via pseudo-bulk subtype assignment across clusters**. **a.** Comparison of subtype probabilities derived from bulk RNA-seq and single-cell RNA-seq in clusters matching bulk RNAseq classification. For each sample, subtype probabilities are shown for bulk RNA-seq (black) and single-cell RNA-seq cluster–based predictions (orange). Points represent probability estimates, and grey lines connect paired bulk and single-cell values from the same sample and cluster. Panels are faceted by predicted cluster subtype, with independent y-axis scaling. This visualization highlights concordance and divergence in subtype probability estimates across data modalities. **b**,**d**, UMAP embedding of single-cell RNA-seq from sample SCPL000059 (**b**) and SCPL000079 (**d**) showing cluster assignments and coloured according to the respective cluster. **c**,**e**, UMAP from **b** and **d** respectively, annotated according to the predicted subtypes per cluster based on TASC analysis of cluster-level pseudo-bulk profiles. Cluster-specific colors match predicted subtype labels. **f**, Bar plots of SingleR predicted cell type proportions for each cluster of patient SCPL000710, aggregated by the predicted subtype assigned to each cluster. Color represent the SingleR predicted cell type, while the proportion is expressed on the y axis within each cluster (x-axis). **g**, same as **f** but for patient SCPL000059 (**b**). **h**, same as **g** but for patient SCPL000079 (d).

#### Supplementary Figure 5.

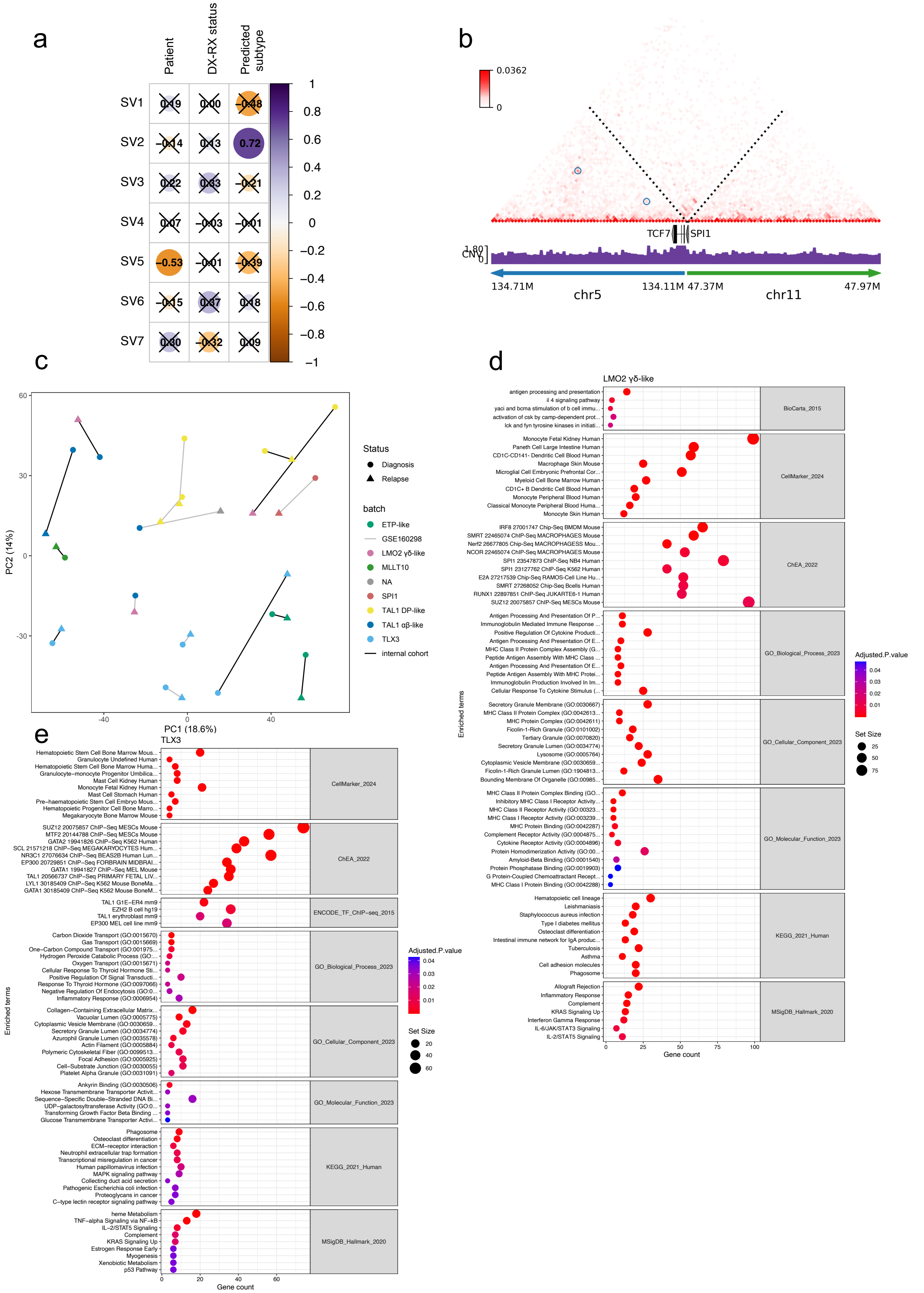

**Differential gene expression analysis and functional enrichment of the selected genes highlight details of subtype specificity of relapse in T-ALL**. **a**, Correlation coefficients between the first seven surrogate variables and key clinical variables (including predicted subtype) are displayed. Dot color and size indicate the direction and magnitude of the correlation (blue = negative, red = positive). A cross marks correlations that did not meet statistical significance ($p \geq 0.05$). **b**, Contact map obtained by HiChIP analysis, with colour scale bar showing distance and cnv-scaled interaction strength at 10 kb resolution mapped onto patient-specific reconstructed linear genome, highlighting the TCF7::SPI1 gene fusion event of patient PAVMUJI at relapse, annotated with hg38 gene annotations. **c**, PCA of the merged and batch corrected dx-rx cohort, showing PC1 vs. PC2 with percentage variance explained. Each point represents a sample colored by subtype. Shape is set to highlight differences between the diagnosis and relapse groups, and each pair is joined by a segment, whose color is representative of the dataset of origin. **d**, Bubble plot of significant EnrichR functional enrichment analysis results (p-adj < 0.05, set size > 3 and OddsRatio > 1) on differentially expressed genes downregulated in relapse ($\log_2$FC > 1.5; p-adj < 0.05) in the TAL1 samples switching to LMO2 γδ-like subtype after relapse. On the x-axis, gene count and dot size are representative of the number of differentially expressed genes within the significant gene set. Color is proportional to adjusted p-value. **e**, Similar to **d**, but on differentially expressed genes upregulated in relapse (-$\log_2$FC > 1.5; p-adj < 0.05) in the TLX3 subtype.

**Supplementary Table 1.**

| | | Sensitivity | Specificity | Pos Pred Value | Neg Pred Value | Precision | Recall | F1 | Prevalence | Detection Rate | Detection Prevalence | Balanced Accuracy |
|---|---|---|---|---|---|---|---|---|---|---|---|---|
| **MVGs** | TLX1 | 0.91 | 0.99 | 0.91 | 0.99 | 0.91 | 0.91 | 0.91 | 0.06 | 0.05 | 0.06 | 0.95 |
| | NKX2-1 | 1.00 | 1.00 | 0.96 | 1.00 | 0.96 | 1.00 | 0.98 | 0.05 | 0.05 | 0.06 | 1.00 |
| | TLX3 | 0.96 | 0.99 | 0.96 | 0.99 | 0.96 | 0.96 | 0.96 | 0.16 | 0.15 | 0.16 | 0.98 |
| | ETP-like | 0.98 | 0.97 | 0.89 | 1.00 | 0.89 | 0.98 | 0.93 | 0.18 | 0.18 | 0.20 | 0.98 |
| | TAL1 DP-like | 0.97 | 0.97 | 0.92 | 0.99 | 0.92 | 0.97 | 0.94 | 0.23 | 0.22 | 0.24 | 0.97 |
| | TAL1 αβ-like | 0.95 | 0.98 | 0.93 | 0.99 | 0.93 | 0.95 | 0.94 | 0.19 | 0.18 | 0.19 | 0.97 |
| | TME-enriched | 0.38 | 0.99 | 0.62 | 0.98 | 0.62 | 0.38 | 0.47 | 0.03 | 0.01 | 0.02 | 0.69 |
| | LMO2 γδ-like | 0.00 | 1.00 | NA | 1.00 | NA | 0.00 | NA | 0.00 | 0.00 | 0.00 | 0.50 |
| | KMT2A | 0.75 | 1.00 | 1.00 | 0.99 | 1.00 | 0.75 | 0.86 | 0.03 | 0.02 | 0.02 | 0.88 |
| | SPI1 | 0.33 | 1.00 | 1.00 | 1.00 | 1.00 | 0.33 | 0.50 | 0.01 | 0.00 | 0.00 | 0.67 |
| | MLLT10 | 0.63 | 0.99 | 0.67 | 0.99 | 0.67 | 0.63 | 0.65 | 0.02 | 0.01 | 0.02 | 0.81 |
| | HOXA9 TCR | 0.86 | 1.00 | 0.92 | 1.00 | 0.92 | 0.86 | 0.89 | 0.02 | 0.01 | 0.02 | 0.93 |
| | BCL11B | 0.67 | 1.00 | 0.89 | 1.00 | 0.89 | 0.67 | 0.76 | 0.01 | 0.01 | 0.01 | 0.83 |
| | STAG2/LMO2 | 0.20 | 1.00 | 0.50 | 1.00 | 0.50 | 0.20 | 0.29 | 0.01 | 0.00 | 0.00 | 0.60 |
| | NKX2-5 | 0.00 | 1.00 | NA | 1.00 | NA | 0.00 | NA | 0.00 | 0.00 | 0.00 | 0.50 |
| **DEGs** | TLX1 | 0.85 | 1.00 | 0.93 | 0.99 | 0.93 | 0.85 | 0.89 | 0.06 | 0.05 | 0.05 | 0.92 |
| | NKX2-1 | 0.82 | 1.00 | 0.97 | 0.99 | 0.97 | 0.82 | 0.89 | 0.05 | 0.04 | 0.05 | 0.91 |
| | TLX3 | 0.95 | 0.98 | 0.92 | 0.99 | 0.92 | 0.95 | 0.94 | 0.16 | 0.15 | 0.16 | 0.97 |
| | ETP-like | 0.97 | 0.94 | 0.77 | 0.99 | 0.77 | 0.97 | 0.86 | 0.18 | 0.18 | 0.23 | 0.96 |
| | TAL1 DP-like | 0.98 | 0.93 | 0.81 | 0.99 | 0.81 | 0.98 | 0.89 | 0.23 | 0.22 | 0.27 | 0.96 |
| | TAL1 αβ-like | 0.90 | 0.99 | 0.95 | 0.98 | 0.95 | 0.90 | 0.93 | 0.19 | 0.17 | 0.18 | 0.94 |
| | TME-enriched | 0.56 | 1.00 | 0.82 | 0.99 | 0.82 | 0.56 | 0.67 | 0.03 | 0.02 | 0.02 | 0.78 |
| | LMO2 γδ-like | 0.00 | 1.00 | NA | 1.00 | NA | 0.00 | NA | 0.00 | 0.00 | 0.00 | 0.50 |
| | KMT2A | 0.75 | 1.00 | 0.95 | 0.99 | 0.95 | 0.75 | 0.84 | 0.03 | 0.02 | 0.02 | 0.87 |
| | SPI1 | 0.33 | 1.00 | 1.00 | 0.99 | 1.00 | 0.33 | 0.50 | 0.01 | 0.00 | 0.00 | 0.67 |
| | MLLT10 | 0.05 | 1.00 | 1.00 | 0.98 | 1.00 | 0.05 | 0.09 | 0.02 | 0.00 | 0.00 | 0.52 |
| | HOXA9 TCR | 0.29 | 1.00 | 0.80 | 0.99 | 0.80 | 0.29 | 0.42 | 0.02 | 0.00 | 0.01 | 0.64 |
| | BCL11B | 0.50 | 1.00 | 1.00 | 0.99 | 1.00 | 0.50 | 0.67 | 0.01 | 0.01 | 0.01 | 0.75 |
| | STAG2/LMO2 | 0.00 | 1.00 | NA | 0.99 | NA | 0.00 | NA | 0.01 | 0.00 | 0.00 | 0.50 |
| | NKX2-5 | 0.00 | 1.00 | NA | 1.00 | NA | 0.00 | NA | 0.00 | 0.00 | 0.00 | 0.50 |

Full-class model train set performance and class prevalence

**Supplementary Table 2.**

| **Subtype** | **Youden's maxima threshold** | **Sensitivity** | **Specificity** | **AUC** | **Specificity maximizing threshold** |
|---|---|---|---|---|---|
| **BCL11B** | **0.368** | **1.0000** | **1.0000** | **1.0000** | **0.368** |
| **ETP-like** | **0.377** | **0.9655** | **0.9974** | **0.9995** | **0.278** |
| **HOXA9 TCR** | **0.188** | **1.0000** | **0.9978** | **0.9997** | **0.188** |
| **KMT2A** | **0.300** | **0.9333** | **1.0000** | **0.9993** | **0.124** |
| **LMO2 γδ-like** | **0.089** | **1.0000** | **1.0000** | **1.0000** | **0.089** |
| **MLLT10** | **0.327** | **0.9091** | **1.0000** | **0.9998** | **0.280** |
| **NKX2-1** | **0.182** | **0.9688** | **0.9977** | **0.9996** | **0.161** |
| **SPI1** | **0.386** | **1.0000** | **1.0000** | **1.0000** | **0.386** |
| **TAL1 DP-like** | **0.350** | **0.9450** | **0.9780** | **0.9952** | **0.261** |
| **TAL1 αβ-like** | **0.429** | **0.9385** | **0.9951** | **0.9988** | **0.374** |
| **TLX1** | **0.235** | **1.0000** | **1.0000** | **1.0000** | **0.235** |
| **TLX3** | **0.336** | **0.9655** | **1.0000** | **0.9995** | **0.231** |
| **TME-enriched** | **0.264** | **0.9412** | **0.9890** | **0.9981** | **0.180** |

Testing set probability-AUCs and their related probability threasholds

**Supplementary Table 3**

| Subtype | term | estimate | std.error | statistic | p.value |
|---|---|---|---|---|---|
| BCL11B | batchGSE110633-636 | -0.140 | 0.097 | -1.444 | 0.199 |
| ETP-like | batchSingle-Cell Pediatric Cancer Atlas | -0.035 | 0.034 | -1.048 | 0.297 |
| ETP-like | batchGSE110633-636 | -0.115 | 0.048 | -2.379 | 0.019 |
| KMT2A | batchSingle-Cell Pediatric Cancer Atlas | -0.149 | 0.152 | -0.977 | 0.340 |
| KMT2A | batchGSE110633-636 | -0.057 | 0.124 | -0.459 | 0.651 |
| LMO2 γδ-like | batchSingle-Cell Pediatric Cancer Atlas | -0.117 | 0.084 | -1.390 | 0.198 |
| LMO2 γδ-like | batchGSE110633-636 | -0.133 | 0.050 | -2.694 | 0.025 |
| MLLT10 | batchSingle-Cell Pediatric Cancer Atlas | 0.049 | 0.140 | 0.353 | 0.730 |
| MLLT10 | batchGSE110633-636 | -0.043 | 0.087 | -0.488 | 0.635 |
| TAL1 DP-like | batchSingle-Cell Pediatric Cancer Atlas | -0.202 | 0.191 | -1.060 | 0.291 |
| TAL1 DP-like | batchGSE110633-636 | -0.140 | 0.060 | -2.331 | 0.021 |
| TAL1 αβ-like | batchSingle-Cell Pediatric Cancer Atlas | -0.019 | 0.098 | -0.189 | 0.850 |
| TAL1 αβ-like | batchGSE110633-636 | -0.035 | 0.045 | -0.776 | 0.440 |
| TLX1 | batchGSE110633-636 | -0.101 | 0.037 | -2.718 | 0.011 |
| TLX3 | batchSingle-Cell Pediatric Cancer Atlas | 0.043 | 0.072 | 0.599 | 0.551 |
| TLX3 | batchGSE110633-636 | -0.233 | 0.042 | -5.621 | 0.000 |

Class specific regression analysis on probability distributions calculating the relative impact of batch and gene imputation

**Supplementary Table 4**

| Cell line name | Literature reported subtype | TASC predicted subtype |
|---|---|---|
| JURKAT | TAL1 | TAL1 DP-like |
| RPMI-8402 | TAL1 | TAL1 αβ-like |
| CCRF-CEM | TAL1 | ETP-like |
| MOLT-4 | TAL1 | TAL1 αβ-like |
| HBP-ALL | TLX3 | TLX3 |
| TALL-1 | TAL1 | TAL1 DP-like |
| DND-41 | TLX3 | TLX3 |
| LOUCY | ETP | ETP-like |
| CUTTL3 | ETP | ETP-like |
| CUTTL1 | TAL1 | TAL1 DP-like |

Validation of TASC predictions on 10 T-ALL cell lines

**Supplementary Table 5.**

| Sample name | ETP status | TASC predicted subtype | TALLSorts predicted subtype |
|---|---|---|---|
| GSM7798273 | ETP | ETP-like | HOXA_MLLT10 |
| GSM7798274 | ETP | ETP-like | HOXA_MLLT10 |
| GSM7798275 | ETP | ETP-like | HOXA_MLLT10 |
| GSM7798276 | ETP | ETP-like | HOXA_MLLT10 |
| GSM7798277 | ETP | ETP-like | HOXA_MLLT10/Diverse |
| GSM7798278 | T-ALL | TAL1 DP-like | Unclassified |
| GSM7798279 | T-ALL | TAL1 DP-like | TAL/LMO |
| GSM7798280 | T-ALL | TAL1 DP-like | TAL/LMO |
| GSM7798281 | ETP | ETP-like | Diverse |
| GSM7798282 | T-ALL | TLX3 | TLX3 |
| GSM7798283 | T-ALL | TLX1 | TLX1 |
| GSM7798284 | T-ALL | TLX1 | TLX1 |
| GSM7798285 | T-ALL | TAL1 αβ-like | TAL/LMO |
| GSM7798286 | T-ALL | TAL1 αβ-like | TAL/LMO |
| GSM7798287 | T-ALL | TAL1 DP-like | TAL/LMO |
| GSM7798288 | T-ALL | TLX1 | TLX1 |
| GSM7798289 | T-ALL | HOXA9 TCR | HOXA_MLLT10 |
| GSM7798290 | T-ALL | TLX3 | TLX3 |
| GSM7798291 | T-ALL | TAL1 DP-like | TAL/LMO |
| GSM7798292 | ETP | ETP-like | Diverse/HOXA_MLLT10 |
| GSM7798293 | ETP | ETP-like | Unclassified |
| GSM7798294 | ETP | TME-enriched | TAL/LMO/Diverse |
| GSM7798295 | ETP | ETP-like | Diverse |
| GSM7798296 | T-ALL | ETP-like | HOXA_KMT2A/TAL/LMO2 |
| GSM7798297 | T-ALL | KMT2A | HOXA_KMT2A/TAL/LMO2 |
| GSM7798298 | T-ALL | TAL1 αβ-like | TAL/LMO |
| GSM7798299 | T-ALL | TAL1 αβ-like | TAL/LMO |
| GSM7798300 | ETP | ETP-like | Diverse |
| GSM7798301 | T-ALL | TAL1 DP-like | NKX2 |

Benchmarking of TASC and TALLSorts predictions on 24 primary samples and 5 cell lines from GSE243914

**Supplementary Table 6.**

| Sample name | ETP status | TALLSorts Predictions | TASC predictions |
|---|---|---|---|
| PPTC-ALL-16-R | ALL | NKX2 | NKX2-1 |
| PPTC-ALL-29-R | ALL | TAL/LMO | TAL1 DP-like |
| PPTC-ALL-33-R | ALL | TAL/LMO | TAL1 DP-like |
| PPTC-ALL-39-R | ALL | TAL/LMO | TAL1 DP-like |
| PPTC-ALL-42-R | ALL | TAL/LMO | TAL1 DP-like |
| PPTC-ALL-46-R | ALL | TAL/LMO | TAL1 DP-like |
| PPTC-ALL-97-R | ALL | TAL/LMO | TAL1 DP-like |
| PPTC-ALL-08-R | ALL | TAL/LMO | TAL1 αβ-like |
| PPTC-ALL-121-R | ALL | TAL/LMO | TAL1 αβ-like |
| PPTC-ALL-27-R | ALL | TAL/LMO | TAL1 αβ-like |
| PPTC-ALL-30-R | ALL | TAL/LMO | TAL1 αβ-like |
| PPTC-ALL-31-R | ALL | TAL/LMO | TAL1 αβ-like |
| PPTC-ALL-43-R | ALL | TAL/LMO | TAL1 αβ-like |
| PPTC-ALL-47-R | ALL | TAL/LMO | TAL1 αβ-like |
| PPTC-ALL-80-R | ALL | TAL/LMO | TAL1 αβ-like |
| PPTC-ALL-81-R | ALL | TAL/LMO | TAL1 αβ-like |
| PPTC-ALL-85-R | ALL | TAL/LMO | TAL1 αβ-like |
| PPTC-ALL-32-R | ALL | TLX3 | TLX3 |
| PPTC-ALL-90-R | ALL | TLX3 | TLX3 |
| PPTC-ETP-1-R | ETP | Diverse | ETP-like |
| PPTC-ETP-3-R | ETP | TAL/LMO | ETP-like |
| PPTC-ETP-5-R | ETP | Diverse | ETP-like |
| PPTC-ETP-6-R | ETP | Unclassified | ETP-like |
| PPTC-ETP-2-R | ETP | TAL/LMO | TAL1 αβ-like |
| PPTC-ETP-4-R | ETP | TLX3 | TLX3 |

Validation and benchmarking of TASC and TALLSorts predictions on 25 PDX samples.

**Supplementary Table 7.**

| Sample name | dx predicted class | rx predicted class |
|---|---|---|
| PARLHD | TAL1 DP-like | TAL1 DP-like |
| PASJFJ | TLX3 | TLX3 |
| PATELN | TAL1 αβ-like | LMO2 γδ-like |
| PAUXAI | TAL1 DP-like | LMO2 γδ-like |
| PAVAIR | TAL1 DP-like | TAL1 DP-like |
| PAVMUJ | SPI1 | SPI1 |
| PAWLSN | TAL1 αβ-like | NA |
| PAYSAZ | TLX3 | TLX3 |

Metadata from dx-rx cohort

**Data S1. (separate file)**
Comprehensive list of data sources and resources used in this study
**Data S2. (separate file)**
Full catalogue of differentially expressed genes (DEGs) identified across subtypes
**Data S3. (separate file)**
Performance metrics for each of the three classification models on the testing set by subtype
**Data S4. (separate file)**
pROC AUCs of test sets after label permutation and retraining for MVG and DEG models, along with ground truth AUCs and pValues
**Data S5. (separate file)**
Training and testing set composition, with model predictions and associated probabilities
**Data S6. (separate file)**
Area under the curve (AUC) values from Precision-Recall (PR) analyses evaluating TASC model performance on the testing set
**Data S7. (separate file)**
Prediction probabilities and class calls for 499 B-ALL samples from the negative validation cohort
**Data S8. (separate file)**
Validation set predictions and probabilities, including 85 primary samples from the GSE110633 and GSE110636 cohorts
**Data S9. (separate file)**
TASC subtype predictions for 10 T-ALL cell lines
**Data S10. (separate file)**
Integrated matched bulk and single-cell RNA-seq (scRNA-seq) prediction data
**Data S11. (separate file)**
Significant differentially expressed gene (DGE) results derived from the dx-rx cohort
